# DeepKin: precise estimation of in-depth relatedness and its application in UK Biobank

**DOI:** 10.1101/2024.04.30.591647

**Authors:** Qi-Xin Zhang, Dovini Jayasinghe, Sang Hong Lee, Hai-Ming Xu, Guo-Bo Chen

## Abstract

Accurately estimating relatedness between samples is crucial in genetics and epidemiological analysis. Using genome-wide single nucleotide polymorphisms (SNPs), it is now feasible to measure realized relatedness even in the absence of pedigree. However, the sampling variation in SNP-based measures and factors affecting method-of-moments relatedness estimators have not been fully explored, whilst static cut-off thresholds have traditionally been employed to classify relatedness levels for decades. Here, we introduce the deepKin framework as a moment-based relatedness estimation and inference method that incorporates data-specific cut-off threshold determination. It addresses the limitations of previous moment estimators by leveraging the sampling variance of the estimator to provide statistical inference and classification. Key principles in relatedness estimation and inference are provided, including inferring the critical value required to reject the hypothesis of unrelatedness, which we refer to as the deepest significant relatedness, determining the minimum effective number of markers, and understanding the impact on statistical power. Through simulations, we demonstrate that deepKin accurately infers both unrelated pairs and relatives with the support of sampling variance. We then apply deepKin to two subsets of the UK Biobank dataset. In the 3K Oxford subset, tested with four sets of SNPs, the SNP set with the largest effective number of markers and correspondingly the smallest expected sampling variance exhibits the most powerful inference for distant relatives. In the 430K British White subset, deepKin identifies 212,120 pairs of significant relatives and classifies them into six degrees. Additionally, cross-cohort significant relative ratios among 19 assessment centers located in different cities are geographically correlated, while within-cohort analyses indicate both an increase in close relatedness and a potential increase in diversity from north to south throughout the UK. Overall, deepKin presents a novel framework for accurate relatedness estimation and inference in biobank-scale datasets. For biobank-scale application we have implemented deepKin as an R package, available in the GitHub repository (https://github.com/qixininin/deepKin).

## Introduction

Detecting relationships among samples is fundamental in genetics and epidemiological analysis, particularly in the context of genome-wide association studies (GWAS) and polygenic risk score (PRS). Conventionally, relatedness has been estimated based on pedigrees, which represents the expected level of genetic similarity. However, with the abundance of genome-wide single nucleotide polymorphism (SNP) data, we can now measure realized relatedness that explicitly captures actual relationship. However, SNP data itself introduces complexity due to a variety of genotyping technologies, quality control (QC) procedures, linkage disequilibrium (LD). Consequently, interpreting estimated relatedness based on genome-wide SNPs can be intricate.

There are various methods to estimate genome-wide relatedness, employing either maximum likelihood approaches (Thompson, 1975; Milligan, 2003; Choi *et al*., 2009) or moments-based estimators (VanRaden, 2008; Manichaikul *et al*., 2010; Yang *et al*., 2010; Conomos *et al*., 2016). Moments-based estimators are often preferred despite their lower precision, because they are computationally efficient (Speed and Balding, 2015). While few factors have been studied on influencing genome-wide relatedness estimations, Hill and Weir made an attempt to explore this within the framework of linkage analysis (Hill and Weir, 2011). They have examined the variation for various pairwise relationship as a consequence of Mendelian sampling and linkage. In contrast, current practice on SNP-based measures is more embraced in the framework of population genome-wide association, while their variation has not been explored. The sampling variance of SNP-based measures varies depending on the LD of SNP data and the level of relatedness. For example, the sampling variance of estimate relatedness can be significantly larger when using more correlated SNPs (due to LD), impacting the statistical power to detect related pairs significantly deviated from unrelatedness. Although factors affecting the variation of method-of-moments relatedness estimators have not been fully explored, static cut-offs have been commonly adopted for inferring relatedness, such as kinship coefficients and IBD coefficients (Manichaikul *et al*., 2010; Ramstetter *et al*., 2017). Neglecting the sampling variance in inference can lead to false positives and misclassifications. Applying static cut-off thresholds without considering sampling variance which is data-specific can generate spurious inferences of relatedness that may not be significantly different from unrelatedness, leading to false positives (Ramstetter *et al*., 2017).

In this study, we present a novel moment-based framework for genome-wide relatedness inference, which is called deepKin. Distinguishing significantly from previous moment estimators, such as KING, deepKin offers advanced capability in relatedness inference, supported by the following statistical features. One remarkable feature is its ability to evaluate and provide the sampling variance of estimated relatedness. By leveraging its asymptotic distribution, deepKin facilitates the computation of *p*-values for each relatedness pair, enabling the assessment of significant deviations from unrelatedness. Furthermore, key principles are emerging when performing relatedness estimation and relatedness inference. I) deepKin determines the critical value that separates significant relatedness estimation from insignificant ones based on sampling variance, which we refer to as deepest significant relatedness; II) it identifies the minimum effective number of markers required for detecting the target degree of relatives to be significantly different from unrelated pairs; and III) Given the target degree of relatedness, it provides the amount of statistical power to be improved or compromised. We verified the performance of deepKin through simulations and demonstrated its effectiveness using the UK Biobank dataset. We also implemented deepKin estimation and the new inference framework in an R package named “deepKin”, which is available in the GitHub repository (https://github.com/qixininin/deepKin).

## Materials and Methods

One of the attempts of the study is to explore the sampling variance for moment-based genetic relatedness. We propose the deepKin estimator, which resembles the KING’s original estimator and differs in the choice of genotype scaling factors. Two forms of genetic relationship matrix (GRM) can be found in Speed and Balding’s review (see Eq 8 and Eq 9 in their review) (Speed and Balding, 2015) or in VanRaden (VanRaden, 2008), which are 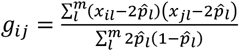 and 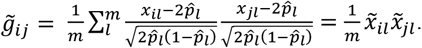. Here, *x*_*il*_ and *x*_*ji*_ are the genotypic values (0, 1, or 2, according to the number of reference alleles) at *l* locus for individuals *i* and *j*, while 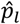 is the allele frequency at *l*-th locus and *m* is the total number of variants.

### KING’s estimator

The original kinship estimator of KING is (Equation 5 in their original publication (Manichaikul *et al*., 2010)

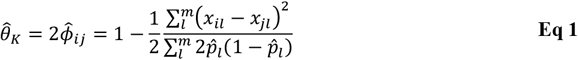

Here the definition of 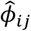 is the same as that in KING’s paper but the expectation of KING’s estimator is only half of the relatedness *θ*. Therefore, in the following text, we tend to use 2 times of the KING’s estimator 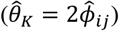 so as to make a clearer comparison (**Table 1**). *θ* has the expected values of 1, 0.5, 0.25, and 0.125 for the zero- (monozygotic twin), first- (full sib, or parent-offspring), second- (half sib, or grandparent-grandchild), and third-degree (first cousin, or great grandparent-great grandchild) relatives, respectively. However, measured through genome-wide similarity, the realized value of *θ* could cover any value between 0 and 1. Note that, the sampling variance of *θ*_1_ has not been explored since it was proposed and the absence of sampling variance hinders the feasibility of conducting precise statistical tests in the statistical framework.

**Table 1.**
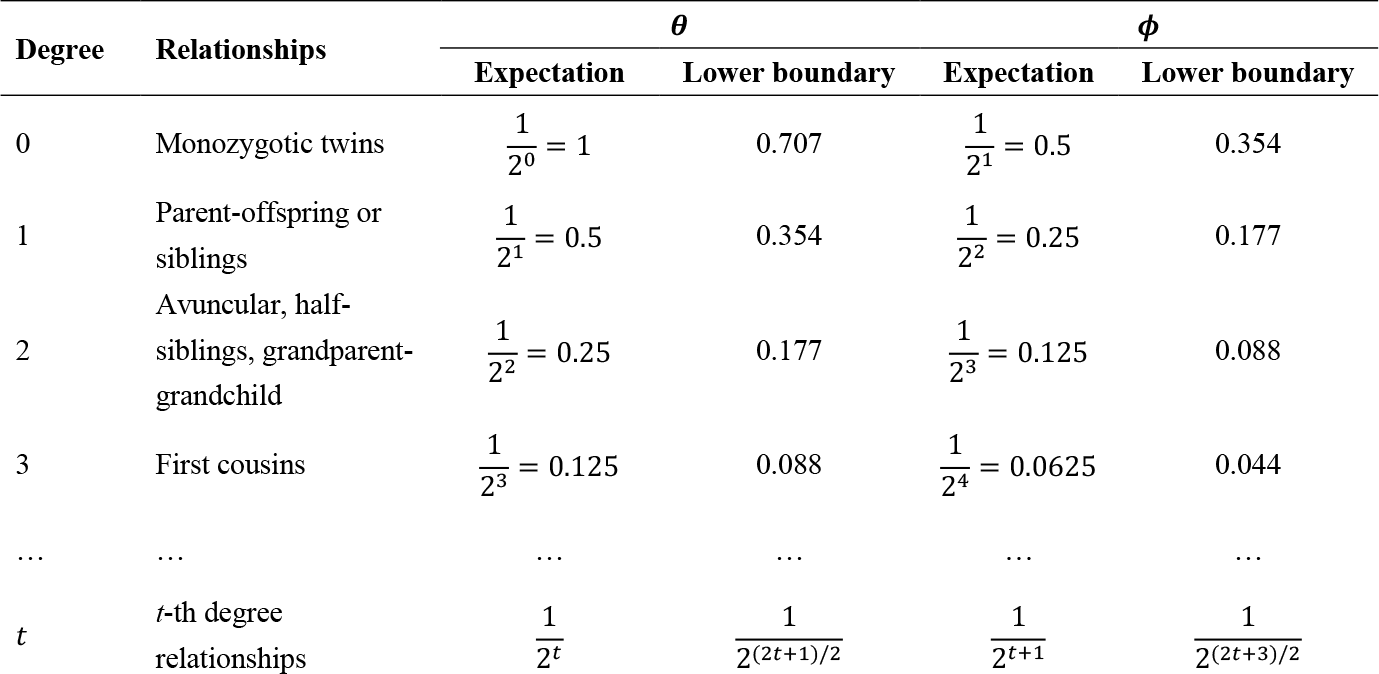
Degrees of relatedness with expected relatedness scores (*θ*) and kinship coefficients (*ϕ*). Includes example relationships for each degree and the corresponding lower boundaries from KING. Table does not include all possible relationship types for these degrees of relatedness. Note that, this table shows relatedness up to the *t*-th degree. Actually, in KING’s original publication, they only considered relatedness up to third-degree, and any relatedness score <0.088 is considered as unrelated.

| Degree | Relationships | $\theta$ | | $\phi$ | |
| --- | --- | --- | --- | --- | --- |
|  |  | Expectation | Lower boundary | Expectation | Lower boundary |
| 0 | Monozygotic twins | $\frac{1}{2^0} = 1$ | 0.707 | $\frac{1}{2^1} = 0.5$ | 0.354 |
| 1 | Parent-offspring or siblings | $\frac{1}{2^1} = 0.5$ | 0.354 | $\frac{1}{2^2} = 0.25$ | 0.177 |
| 2 | Avuncular, half-siblings, grandparent-grandchild | $\frac{1}{2^2} = 0.25$ | 0.177 | $\frac{1}{2^3} = 0.125$ | 0.088 |
| 3 | First cousins | $\frac{1}{2^3} = 0.125$ | 0.088 | $\frac{1}{2^4} = 0.0625$ | 0.044 |
| ... | ... | ... | ... | ... | ... |
| $t$ | $t$-th degree relationships | $\frac{1}{2^t}$ | $\frac{1}{2^{(2t+1)/2}}$ | $\frac{1}{2^{t+1}}$ | $\frac{1}{2^{(2t+3)/2}}$ |

### deepKin estimator

Inspired by the construction of two GRMs with different scaling factors and in contrast to **Eq 1**, the deepKin estimator 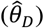 is constructed as,

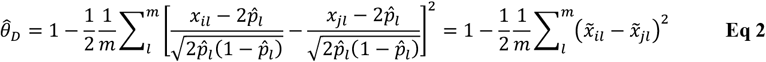

After some rearrangement, **Error! Reference source not found**. can be decomposed into three parts, which are 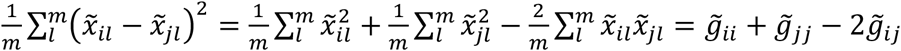. Both 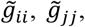, and 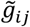 are matrix elements of the GRM, which has the computational cost of 𝒪(*n*^2^*m*). This decomposition gives the relationship between GRM and deepKin that 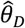 for any pair of individuals can be quickly realized through addition of these three GRM elements. The computational cost of deepKin is consequently 𝒪(*n*^2^*m*), which is identical to the cost of constructing GRM.

We first give the expectation and variance of deepKin under the assumption of a binomial distribution at one locus, namely the “single locus model” (see **Appendix I**). The expectation and variance of deepKin for a marker locus 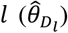 are

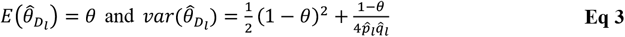

However, in practice we have many variants that are often in LD, therefore, we derive the expectation and variance of deepKin under the “multiple locus model” (see **Appendix II**). We assume that the aggregated standardized genotypes follow the normal distribution and have the expectation of 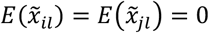 and variance of 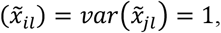, which meets the requirement of Isserlis’s theorem (Isserlis, 1918). After recursively applying Isserlis’s theorem, we are able to derive the expectation and asymptotic variance of 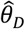, which are,

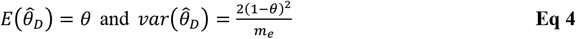

The sampling variance of 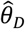 between two individuals is subject to the true relatedness (*θ*, lessened sampling variance for more related samples) and the global LD between variants 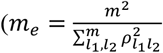 is the effective number of markers, in which 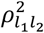 is squared correlation between loci *l*_1_ and *l*_2_ (see more discussion on *m*_*e*_ below). 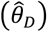 depends on the underlying normal distribution for the aggregated variants and may be disrupted by low-frequency variants, therefore, we restrict MAF to at least 0.05. For details of deriving the expectations and variances under “single locus model” and “multiple loci model” please refer to **Appendix I and II**. The “single locus model” is exact but is limited to independent variants, while the “multiple loci model” is asymptotic but takes into account the global LD between *m* variants. Therefore, the multiple loci model is preferred in the real data analysis and has been adopted to perform subsequent relatedness inferences. Considering the context of **Eq 3** and **Eq 4**, a negative 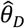 can be attributed to sampling variance, but it may also imply possible diversity among the samples.

### Relatedness inference: deepest significant estimation

The deepKin’s framework provides statistical inference for the estimated relatedness score. To test whether a pair of individuals is related, the null distribution of unrelated pairs is 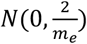 given by **Eq 4**. Therefore, the z score and corresponding *p*-value on 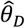 are 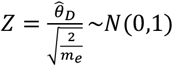 and *p* = 1 − *ϕ*(*Z*). *ϕ* is the standard normal cumulative distribution function. If we assume a significant level of *α*, the critical value 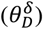 is

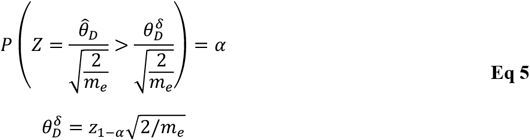

The critical value is considered as the deepest significant estimation and 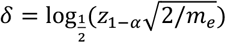 will be the deepest significant degree by log-transformation from relatedness to relationship degree.

### Relatedness inference: classification

Although the relatedness score can be continuous, we classify them into discrete classes for easier access, as often done in real data analysis (**Table 1**). Since **Eq 4** gives the sampling variance for any relatedness, we can make a concrete inference for any observed significant relatedness between *t* and *t* + 1 degree using hypothesis testing. The null distributions of the relatedness estimations at *t* and *t* + 1 degrees are 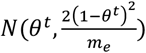 and 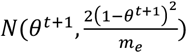 respectively. According to definition, we have 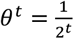 and 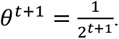. The classification of *x* degree to *t* or *t* + 1 degree is performed based on two *Z*-test statistics. For hypothesis *H*_0_: *θ*^*x*^ = *θ*^*t*^, *H*_1_: *θ*^*x*^ < *θ*^*t*^, we can calculate its z score and *p*-value of rejecting *H*_0_ as

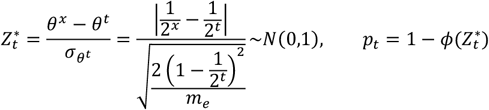

For hypothesis *H*_0_: *θ*^*x*^ = *θ*^*t*+1^, *H*_1_ *θ*^*x*^ > *θ*^*t*+1^, we can calculate its z score and *p*-value of rejecting *H*_0_ as

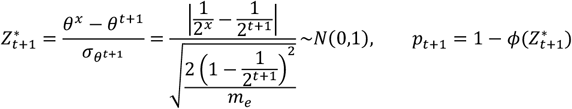

By comparing these two *p*-values, we are able to infer the classification for any observed *θ*^*x*^. Intuitively, there exists a specific point of relatedness where the two *p*-values are equal. At this crossover point for *p*-values, the relatedness value can be utilized as a new boundary for direct classification (**Appendix III**).

### Derived guidelines from deepKin

Two parameters are involved in the estimation of relatedness with deepKin: the sample size (*n*) and the number of markers (*m*). However, it is actually *m*_*e*_ rather than *m* that acts as an indicative parameter in relatedness inference, affecting the performance of deepKin. Making relatedness inference of *n*(*n* − 1)/2 pairs of samples is a decision-making process. Our subsequent presentation follows the framework of power calculation, which aims to consider Type-I (*α*) and Type-II (*β*) error rates and offer preliminary yet practical guidelines that can be applied.

#### Guideline I: thrifty choice for markers conditional on a target degree

Based on the expectation and variance of deepKin estimator under “multiple locus model”, we can estimate the minimum *m*_*e*_ that is required for detecting the target degree (*t*) of relatedness 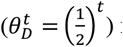 from unrelated pairs, which is related to Type-I and Type-II error rates (*α* and *β*),

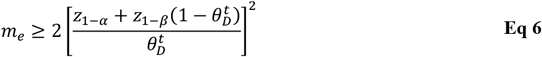

Here, *z*_1−*α*_ and *z*_1−*β*_ are the quantiles from the standard normal distribution. For more details see **Appendix IV**. In particular, we set *α* under experiment-wise control after Bonferroni correction, the corresponding *α* is upon the total comparisons *N*. We give a concrete numerical example for a simple illustration. Suppose that we have 500,000 individuals, which generates *N* ≈ 1.25*e*11 pairs of comparisons. We wanted to detect relatives up to the third-degree from unrelated pairs at Type I error rate of *α* = 0.05/*N* (*z*_1-*α*_ = 7.161) and Type II error rate of *β*=0.1 (*z*_1−*β*_ = 2.326), and the number of *m*_*e*_ based on **Eq 6** should be no less than 8,780. Then a thrifty choice of a variant set can greatly reduce the financial cost of genotyping and computation. This gives the first guideline that select only a subset of the genetic variants with the required *m*_*e*_ is sufficient for detecting certain degree of relatedness in *N* comparisons. Consequently, *m*_*e*_/*m* reflects the balance between statistical power and computational cost though economical choice for markers.

#### Guideline II: the statistical power conditional on target degree and me

As often *α* is fixed in a study – such as after Bonferroni correction, we are able to determine how much of the power (*π*) could be compromised or improved for any target relatedness 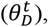,

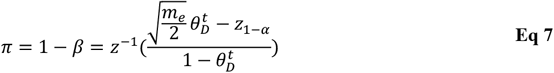

For the same example above, if the target degree is 5, the power increases to 0.925; and if the target degree is 6, the power is rather reduced to 0.002. To increase statistic power, an applicable way is to increase the effective number of markers. When the effective number of markers increases, the power to detect the target relatedness increases (**Figure S1**). For more real data examples please refer to **Table S1**.

### On the effective number of markers (*m*_*e*_)

The above analysis relies on the effective number of markers *m*_*e*_, which is defined as 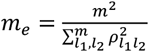 so as to describe the average LD between genome-wide variants (Chen, 2014; Visscher *et al*., 2014; Zhou, 2017). Intuitive *m*_*e*_ can be directly calculated through calculating all squared correlations between variant pairs (such as –r2 command in PLINK), but the computational cost is 𝒪(*nm*^2^) and it soon becomes unaffordable when *n* and especially, *m*, is large. However, estimating *m*_*e*_ by direct calculation can be computationally substantial, whilst *m*_*e*_ is needed as a hyperparameter in determining the potential of the data such as characterized by **Eq 5-7**. Here, we propose a pair of estimators for *m*_*e*_.

#### Estimator I: GRM-based estimator

We first introduce the GRM-based estimator below

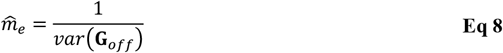

in which **G**_*off*_ refers to the off-diagonal elements of GRM, namely GRM-based estimator in the following context (Chen, 2014). 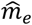 estimated from **Eq 8** is asymptotically normal with its sampling variance of 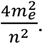. The computational cost is 𝒪(*n*^2^*m*), which is often smaller than 𝒪(*nm*^2^). A more detailed investigation on the relationship between genome-wide LD and GRM-based *m*_*e*_ estimator can be found in Huang *et al*. (Huang *et al*., 2023).

#### Estimator II: randomization-based estimator

The second estimator is based on a randomization algorithm, which reduces the computational cost of estimating *m*_*e*_ from 𝒪(*n*^2^*m*) to 𝒪(*nmB*). Here, *B* is the number of iterations and is practically sufficient if *B* > 100. The randomization method uses an *n* × *B* matrix **S**, whose element is randomly sampled from the standard normal distribution. By calculating a statistic 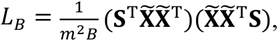, where 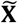 is the standardized genotype matrix, the empirical randomization-based estimator is then 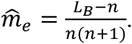. Its asymptotic sampling variance is 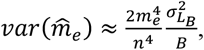, in which 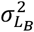 can be estimated from the *B* rounds of iteration for the estimation of *L*_*B*_. The randomized estimation is inspired by randomized estimation of heritability (Wu and Sankararaman, 2018).

In the simulations below, *m*_*e*_ was estimated using GRM-based estimator. Even though GRM will be calculated during deepKin estimation and consequently *m*_*e*_ could be estimated, the randomization-based estimator still has its advantages of a faster approximation thus a faster evaluation of the guidelines, which promises a faster decision on SNP set selection.

### Simulation

#### Variance validation

We first validated the single locus model (**Eq 3**). *n* = 2,000 pairs of individuals with relatedness score *θ* were simulated, and *θ*=0.5, 0.25, 0.125, and 0, respectively. 5,000 equal-frequency markers were simulated, and the frequency were 0.01, 0.05, 0.10, 0.15, 0.20, 0.25, 0.30, and 0.35, respectively. Both expected and observed variances were evaluated among 2,000 pairs of individuals. Standard deviation of each variance estimation was evaluated through ten repeats.

To validate **Eq 4** based on multiple loci model, we simulated the same *n* = 2,000 pairs of individuals using 5,000 markers but whose MAFs were sampled from a uniform distribution, which were *U*(0.01, 0.5), *U*(0.05, 0.5), *U*(0.10, 0.5), *U*(0.15, 0.5), *U*(0.20, 0.5), *U*(0.25, 0.5), *U*(0.30, 0.5), and *U*(0.35, 0.5), respectively. We used *D*′ to represent the LD, which was sampled from a uniform distribution *U*(0.1, 0.2) and *U*(0.5, 0.8) to represent low and high LD conditions. *m*_*e*_ used to calculate the asymptotic variance was estimated from GRM-based estimator. Ten repeats were used to achieve the standard deviation of each variance estimation.

#### Data-based threshold determination

We showed the difference in the inference frameworks between KING and deepKin in simulation. Only unrelated individuals (*n* = 2,000) were simulated based on different number of markers (*m* = 1,000, 5,000, 10,000, and 50,000). The genotype simulation followed the LD scenario as described above. MAF was sampled from a uniform distribution *U*(0.05, 0.5) and *D*′ was sampled from a uniform distribution *U*(0.1, 0.2), and correspondingly *m*_*e*_ = 978, 4,896, 9,782, and 48,967 as estimated by the GRM-based estimator. We applied the KING’s inference criteria of 0.707 (the geometric mean of 1 and 0.5, **Table 1**) for monozygotic twins and duplicate samples, 0.354, 0.177, 0.088, and 0.044 for first-, second-, third-, and fourth-degree of relatedness, respectively. We calculated the deepest significant relatedness estimation (*θ*^*δ*^) from **Eq 5** based on *m*_*e*_ at the significant level of *α*=0.05/1,999,000. This enables data-specific thresholds for relatedness inference.

#### Relatedness inference based on p-values

To validate the role of *p*-values in inferring relatedness, we simulated various related pairs in two cohorts. We simulated 200 individuals each for cohort 1 and cohort 2 (*n*_1_ = *n*_2_ = 200). Between cohort 1 and cohort 2, we generated 10 pairs of related samples up to the fourth-degree. MAF was sampled from a uniform distribution *U*(0.05, 0.5) and *D*′ was sampled from a uniform distribution *U*(0.1, 0.2). The numbers of markers were *m*=133, 687, 3,086, 13,050, and 53,643, corresponding to identical, first-, second-, third-, and fourth-degree relatives, respectively. Taking fourth-degree as an example, the number of markers was determined by the following procedures. the minimum number of *m*_*e*_ based on **Eq 6** for fourth-degree was 17,881 at Type I error rate of *α*=0.05/40,000 and Type II error rate of *β*=0.1. To make sure that the actual *m*_*e*_ of simulated markers meets the requirement, we simulated *m* markers that are three times of the minimum *m*_*e*_, *m* = 3*m*_*e*_ = 53,643. The actual *m*_*e*_ of 53,643 markers were 18,612. All sets of selected markers met the requirement of the target minimum number of *m*_*e*_. *m*_*e*_ was estimated from GRM-based estimator. The deepest significant relatedness estimations were calculated according to *m*_*e*_ based on **Eq 5** at a significant level of 0.05/40,000.

### UK Biobank application

#### Oxford demo for the proof of principle

We drew 3,000 UK Biobank (UKB) participants from Oxford and analyzed four sets of SNPs, each representing a different effective number of markers. The first set had 693,666 imputation loci after QC, and the distribution of MAF was as shown in **Figure S2**. The inclusion criteria for autosome variants were: i) minor allele frequency (MAF) > 0.05; ii) Hardy-Weinberg equilibrium (HWE) test *p*-value > 1e-7; iii) no locus missingness. The second SNP set had the same inclusion criteria as SNP set 1, but the MAF threshold was increased from 0.05 to 0.2 and resulted in 298,221 SNPs. The third SNP set excluded variants that had high population differentiation in SNP Set 2, which remained 237,642 variants and were often called as non-ancestry informative markers (non-AIM) (Chen *et al*., 2016). The fourth SNP set was performed LD pruning on SNP set 2 (*r*^2^<0.1 in a 50-variant window and a 5-variant count to shift the window), which reduced the number of variants to 36,425.

We estimated *m*_*e*_ for each of the four SNP sets using the randomization method. To further describe how the choice of the SNP sets could result in the fluctuation of *m*_*e*_, we examined a total of 13 conditions, 6 of which were randomly selected markers with different sizes (*m* = 5,000, 10,000, 50,000, 100,000, 150,000, and 200,000) and 7 of which had different pruning thresholds (*r*^2^< 0.05, 0.1, 0.2, 0.3, 0.4, 0.5, and 0.8 in a 50-variant window and a 5-variant count to shift the window). All conditions were performed on SNP set 2.

#### Relatedness in UKB white British ancestry subset

We considered a subset of 427,287 participants with self-reported white British ancestry from 19 assessment centers with a sample size greater than 5,000. Leeds had the largest sample size of 39,707, while Bury had the smallest sample size of 7,701. 72,016 imputation SNPs remained after QC (**Figure S3**). The inclusion criteria for autosome variants were: i) MAF > 0.05; ii) HWE test *p*-value > 1e-7; iii) no locus missingness; iv) *r*^2^<0.1 in a 50-variant window and a 5-variant count to shift the window. To explore the geographical connection between relatives, we included the grid co-ordinates for 19 assessment centers, which were downloaded from the UKB website (https://biobank.ndph.ox.ac.uk/ukb/refer.cgi?id=11002).

## Results

### Simulation results

We first validated the variance derived from deepKin in two models, the “single locus model” and the “multiple loci model”. By simulating paired individuals with relatedness up to third-degree and calculating their relatedness estimation using KING-homo and deepKin, we presented their observed variances, together with the expected variances derived under the “single locus model” (**Figure 1 A-D**). The observed variances of KING-homo and deepKin did not differ significantly from the expected one at all degrees and all MAF scenarios. When a more generalized context was introduced – MAFs were randomly assigned and different conditions of LD were considered, the observed variances resembled the expected variances derived under the “multiple loci model” at all degrees of relatedness and fluctuated at different MAF scenarios (**Figure 1 E-L**). The two observed variances of KING-homo and deepKin showed no significant difference when the lower boundary of MAF was above 0.1. It was worth noting that, the inclusion of low-frequency variants (MAF<0.05) introduced an increase of sampling variance significantly. With choice of common variants, the expected variance is often conservative to the observed one.

**Figure 1.**
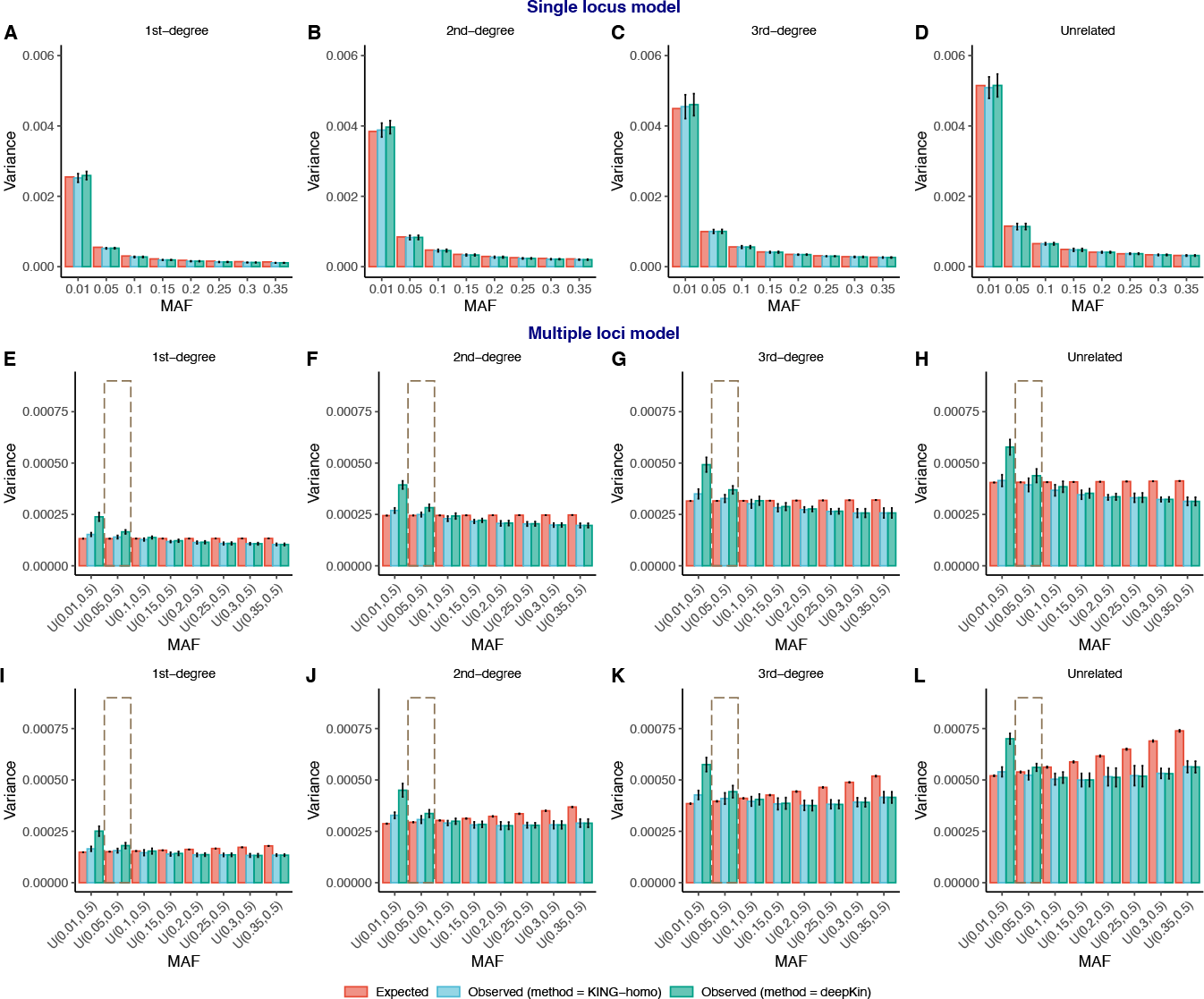
The expected and observed variances under “single locus model” and “multiple loci model” of different MAF scenarios in simulation. (**A-D**) “Single locus model”: 5,000 independent markers of the same MAF (0.01, 0.05, 0.10, 0.15, 0.20, 0.25, 0.30, and 0.35) are simulated. (**E-L**) “Multiple loci model”: 5,000 dependent markers of different MAFs sampled from different uniform distributions are simulated, which are *U*(0.01, 0.5), *U*(0.05, 0.5), *U*(0.10, 0.5), *U*(0.15, 0.5), *U*(0.20, 0.5), *U*(0.25, 0.5), *U*(0.30, 0.5), and *U*(0.35, 0.5). *D*^2^ is used to depict the linkage disequilibrium and is sampled from a uniform distribution *U*(0.1, 0.2) and *U*(0.5, 0.8) to represent low LD (**E-H**) and high LD (**I-L**). A total of 2,000 first-degree, second-degree, third-degree, and unrelated pairs are simulated and evaluated. All scenarios are performed in 10 repeats. 95% confidence intervals are given as error bars.

We then performed simulation to intuitively show the difference in the inference of relatedness between KING and deepKin (**Figure 2**). Only unrelated individuals were simulated. deepKin took into account the distribution of null hypothesis, which was related to the effective number of markers of the data, and provided dynamic cut-offs for relatedness inference where all unrelated pairs were inferred as “insignificant” in all scenarios. The quantile-plots for the *p*-value of deepKin showed that the observed distribution matched with our expectation. However, fixed thresholds of KING could lead to false positives either because there was limited number of markers or the target degree was too distant, for instance, the lower boundary for third-degree (0.088) was within the distribution of unrelated individuals when *m*=1,000 and 5,000, and the lower boundary for fourth-degree (0.044) was within the distribution of unrelated individuals when *m*=1,000, 5,000, and 10,000.

**Figure 2.**
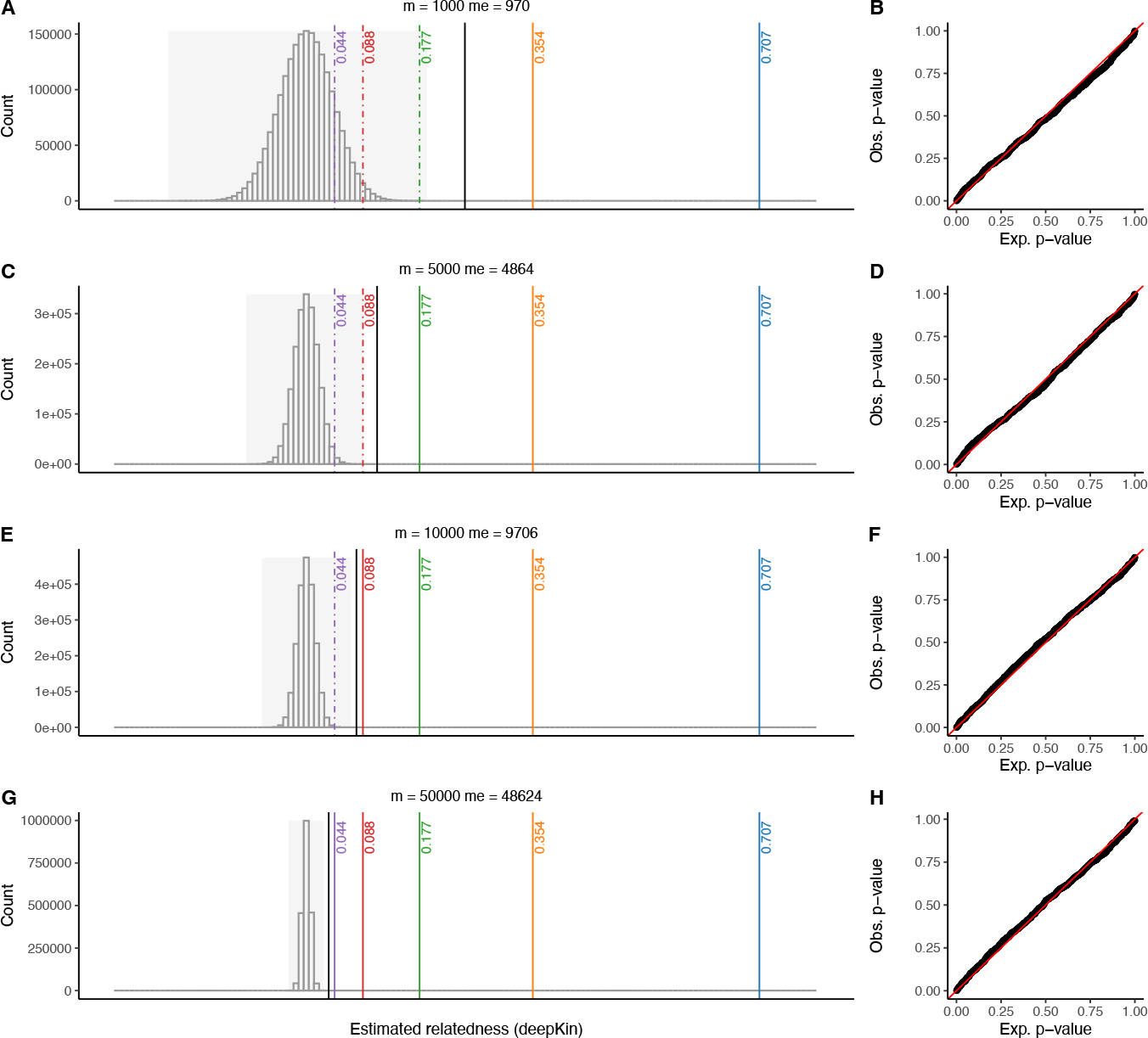
Different ways of relatedness inference between KING and deepKin. (**A, C, E, and G**) Histograms of relatedness score estimations by deepKin on unrelated individual pairs using different number of markers (*m* = 1,000, 5,000, 10,000, and 50,000). Only unrelated individuals (*n*=2,000) were simulated. Line in black indicates the deepest significant relatedness calculated at the significant level of *α*=0.05/1,999,000. Lines in color indicate KING’s lower boundaries of 0.707 (blue), 0.354 (yellow), 0.177 (green), 0.088 (red), and 0.044 (purple) for zero-, first-, second-, third-, and fourth-degree of relatedness. Those lower boundaries that is smaller than deepest relatedness supported by deepKin is plotted in dashed lines. The grey background indicates the observed relatedness range. (**B, D, F, and H**) The quantile-plots for deepKin *p*-values. The *p*-values on x-axis are sampled from the uniform distribution *U*(0, 1).

We also demonstrated the distribution of *p*-values for different degrees of relatives (identical, first-, second-, third-, and fourth-degrees) in simulation (**Figure 3**). The closer the relatives were, the more significant their *p-*values were. It turned out that when the actual size of *m*_*e*_ met the requirement of that for target degree at the experiment-wise Type I error rate of 0.05 and Type II error rate of 0.1 based on guideline I, all target relatives were clearly separated from unrelated pairs based on the significant thresholds.

**Figure 3.**
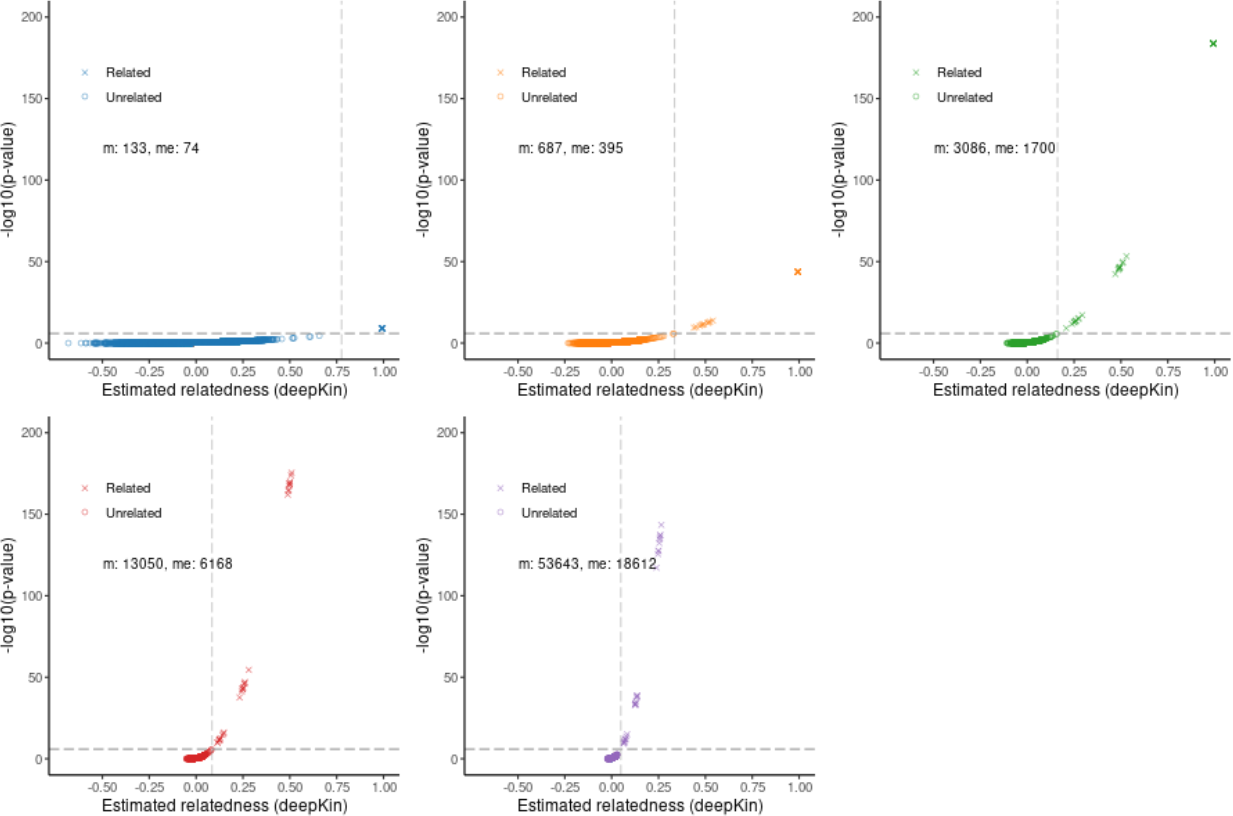
The distribution of *p*-values for different degrees of relatives in simulation up to fourth-degree. The plot shows the estimated relatedness scores and the corresponding *p*-values for unrelated (circle) and related (cross) pairs. The horizontal line indicates the significant level of *α*=0.05/40,000. The vertical dashed line indicates the corresponding deepest significant relatedness based on **Eq 5**.

### UKB Oxford demo

We first described how *m*_*e*_ was fluctuated with different choices of markers in the Oxford demo (**Figure S4**). A total of 13 conditions were made up of two procedures, random selection or LD pruning. By randomly selecting variants along the genome, *m*_*e*_ increased sharply at first and then slowly, eventually approximating that of the overall QCed variants. For example, the number of markers and the effective number of markers were *m* = 298,211 and *m*_*e*_ = 7,634 for all SNPs. However, when the threshold of LD pruning was *r*^2^ < 0.4, the number of markers and the effective number of markers were *m* = 91,670 and *m*_*e*_ = 41,000.

Based on these four sets of SNPs, we demonstrated the role of *m*_*e*_ in affecting the sampling variance of relatedness estimation and subsequently relatedness inference using 3,000 Oxford samples of white British ancestry. deepKin took *m*_*e*_ into consideration and calculated the deepest significant relatedness supported by each SNP set (**Table 2**). The degrees of deepest significant relatedness supported by SNP sets 1-3 were closer than the fourth degree, while SNP set 4, which had the largest size of *m*_*e*_, harbored the deepest significant degree up to 4.409. Therefore, SNP set 4 discovered a total number of 57 significant relatedness, while SNP set 1-3 discovered 37, 32, and 38 significant relatedness (see **Data S1**). We showed the expected power and classification *p*-values using these four sets of SNPs (**Figure 4A and 4B**). SNP set 4, which had the largest *m*_*e*_ and the smallest expected sampling variance, offered the most powerful inference for distant relatives and was the most reliable one in relatedness classification. Taking SNP set 2 and 4 as examples, deepKin estimations were very consistent to the “KING-robust” estimations at positive values (**Figure 4C and 4D**). No additional relatives were found for deepKin under two different SNP sets. However, casual usage of KING’s relative inference cut-offs, which were constant values regardless of the changing of SNP sets, might lead to substantial difference (**Table 2**).

**Table 2.**
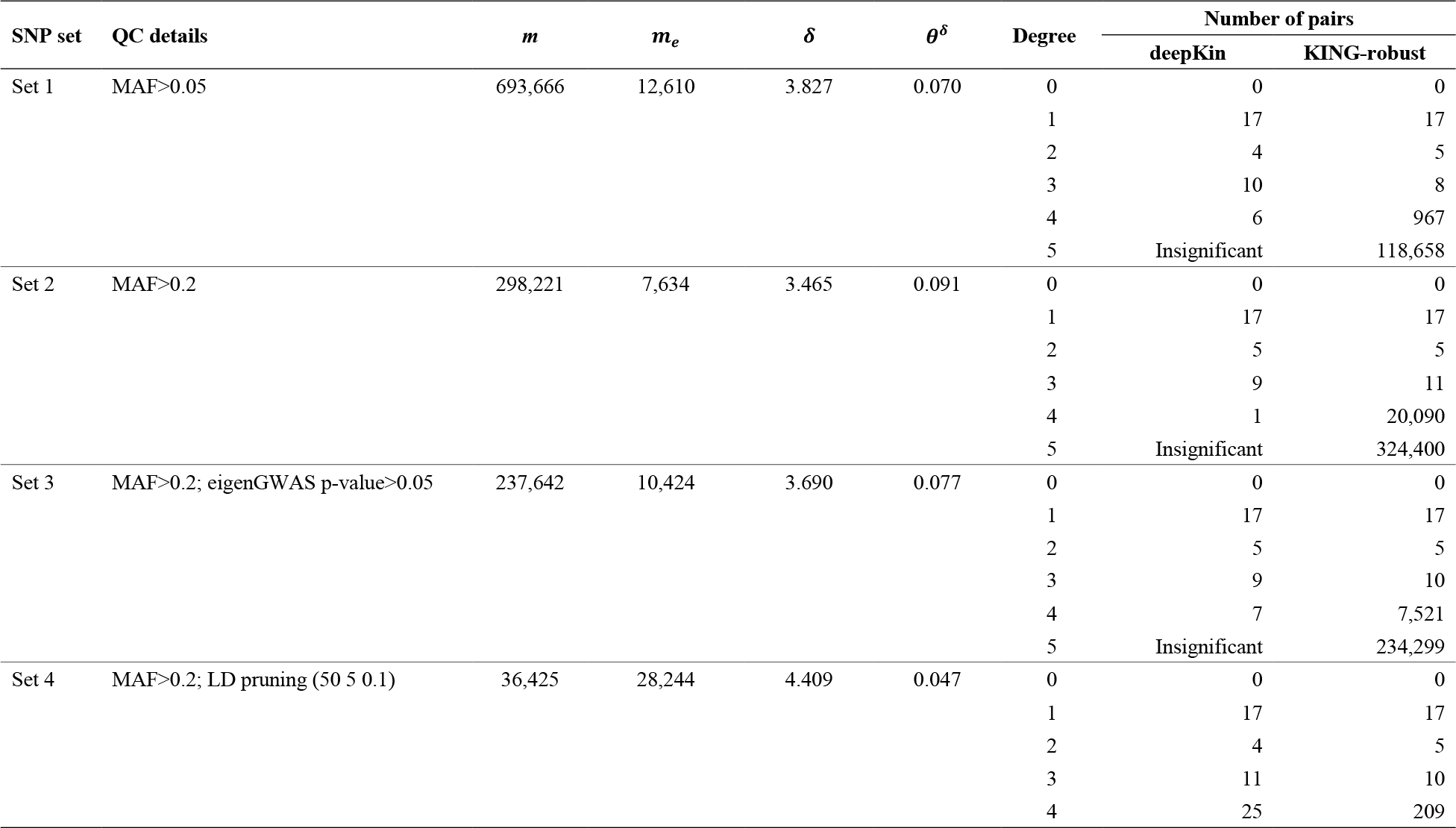
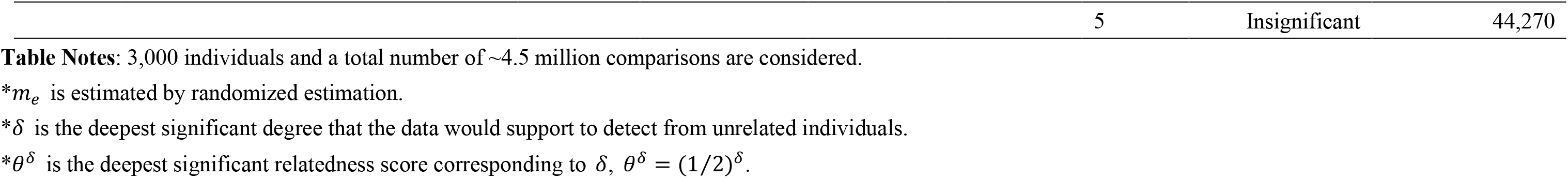
QC details for four SNP sets and the numbers of pairs assigned to each degree of relationship based on deepKin and KING framework in the Oxford demo.

**Figure 4.**
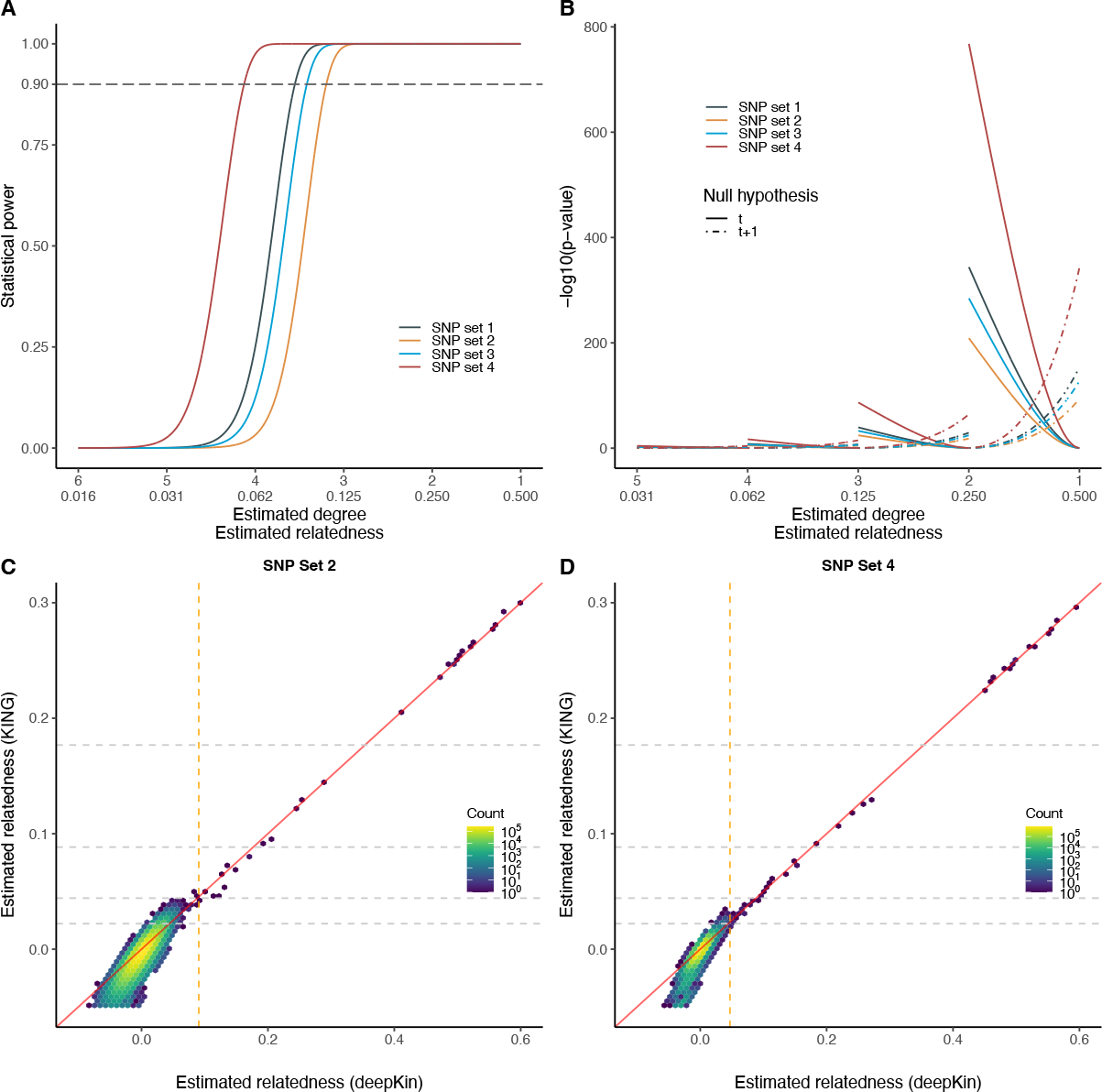
Oxford demonstration. (**A**) The expected power of detecting different relatedness using four SNP sets. The dashed grey line indicates 90% power. Suppose Type I error rate of *α* = 0.05/*N* and Type II error rate of *β*=0.1, where *N* = 4,498,500. (**B**) The expected *p*-values for relatedness classification using four SNP sets. *p*-values under two null hypotheses on adjacent *t* (solid curves) and *t* + 1 (dashed curves) degree are plotted. (**C** and **D**) Scatter plots of the estimated relatedness by KING-robust and deepKin for all individual pairs in Oxford 3K demo using SNP set 2 (**C**) and SNP set 4 (**D**). Colors indicate the number of pairs that fall within the range of each hexagonal bin. Dashed grey lines on y-axis indicate KING’s cut-offs while the dashed orange line indicates deepKin’s critical value at significant level of *α* = 0.05/*N*.

To further investigate relatedness classification, the numbers of pairs assigned to each degree of relationship based on deepKin and KING were compared (**Table 2**). The numbers of relative pairs were quite consistent among the four SNP sets for identical and first-degree pairs (0 and 17 respectively) using deepKin or KING. The numbers of relative pairs were also quite consistent for second- and third-degree, 4 or 5 for second-degree, and 8 to 11 for third-degree. However, for fourth-degree relatives, the number of relative pairs discovered by KING inflated and fluctuated dramatically among the four SNP sets, which were 2,769, 20,090, and 7,521 using SNP set 1-3 and 209 using SNP set 4. As deepKin only performed classification on those significant signals, no abnormal inflation of distant relatives should be observed. Simple inference based on static criteria without considering the sampling variance could introduce unexpected false positives, at least for distant relatives who might be slightly related but were not fully supported by the data. The calculation of deepest significant relatedness by deepKin offered safe examinations for each data set, and supported further relatedness classification.

As GRM was often used for sample QC in GWAS, we also investigated the number of pairs that exceed the normal cut-offs for GRM (**Table S2**). Using different SNP sets, the numbers of excluded relative pairs ranged from 60 to 2,381 and from 655 to 72,032 using GRM’s fixed cut-offs *θ*>0.05 and *θ*>0.025, respectively. These results provided a comprehensive understanding of how deepKin interacted with both the data itself and user-specified QC.

As low-frequency variants might breach the asymptotic sampling variance for deepKin (**Eq 4**), we examined the sampling variance of deepKin estimations in the Oxford demo when using different MAF thresholds (0.01, 0.05, 0.10, 0.15, 0.20, 0.25, 0.30, and 0.35) (**Figure S5**). *m* was the number of variants that remained after applying MAF thresholds and *m*_*e*_ was the effective number of markers calculated from the randomization method. There was no clear gap between the observed histogram and the asymptotic normal distribution with *σ*^2^ = 2/*m*_*e*_ when MAF thresholds were above 0.05. We strongly suggest that low-frequency variants should be removed during quality control.

### UKB white British ancestry subset

We then applied deepKin to the entire subset of participants with white British ancestry in UKB, which had *n*=427,287 and were from 19 assessment centers. After basic QC and LD pruning, the *m*_*e*_ of the 72,016 markers was 56,945. We used deepKin to estimate nearly 𝒩 ≈ 9.13*e*10 pairs, which took approximate 193 hours with 48 threads of two Intel(R) Xeon(R) CPU E5-2690 v3 @ 2.60GHz. A total of 232,552 (54.4%) UKB participants with white British ancestry were inferred to be related to at least one other person in the subset at the significant level of 0.05/𝒩, and formed a total of 212,120 statistically significant related pairs (**Figure 5A**). Based on **Eq 5**, this SNP set held the deepest significant degree of *δ*=4.567 (95% CI: 4.565∼4.569). These 212,120 significantly relatedness estimations were classified into six degrees (**Table S3**): 162 pairs of identical pairs/monozygotic twins, 25,699 pairs of first-degree relatives, 9,455 pairs of second-degree relatives, and 53,221 pairs of third-degree relatives. Dividing by the total number of comparisons 𝒩, these forms were equivalent, but different numbers, to the relative components as reported in the original UKB report (Bycroft *et al*., 2018). Besides, a total of 129,930 (30.4%) participants with white British ancestry were related up to at least third-degree, where this ratio for all participants was 30.3% in Bycroft *et al*. (Bycroft *et al*., 2018). Moreover, since the deepest significant degree was deeper than third-degree, we also reported 91,977 and 31,606 pairs of fourth- and fifth-degree relatives. Pairs of related individuals within the UKB white British subset formed networks of related individuals. While in most cases these were networks of size two or three, there were also many groups of size four or even larger in the subset (**Figure 5B**). If we only considered related individuals up to third-degree, then the largest group size was reduced to 10 (data not shown).

**Figure 5.**
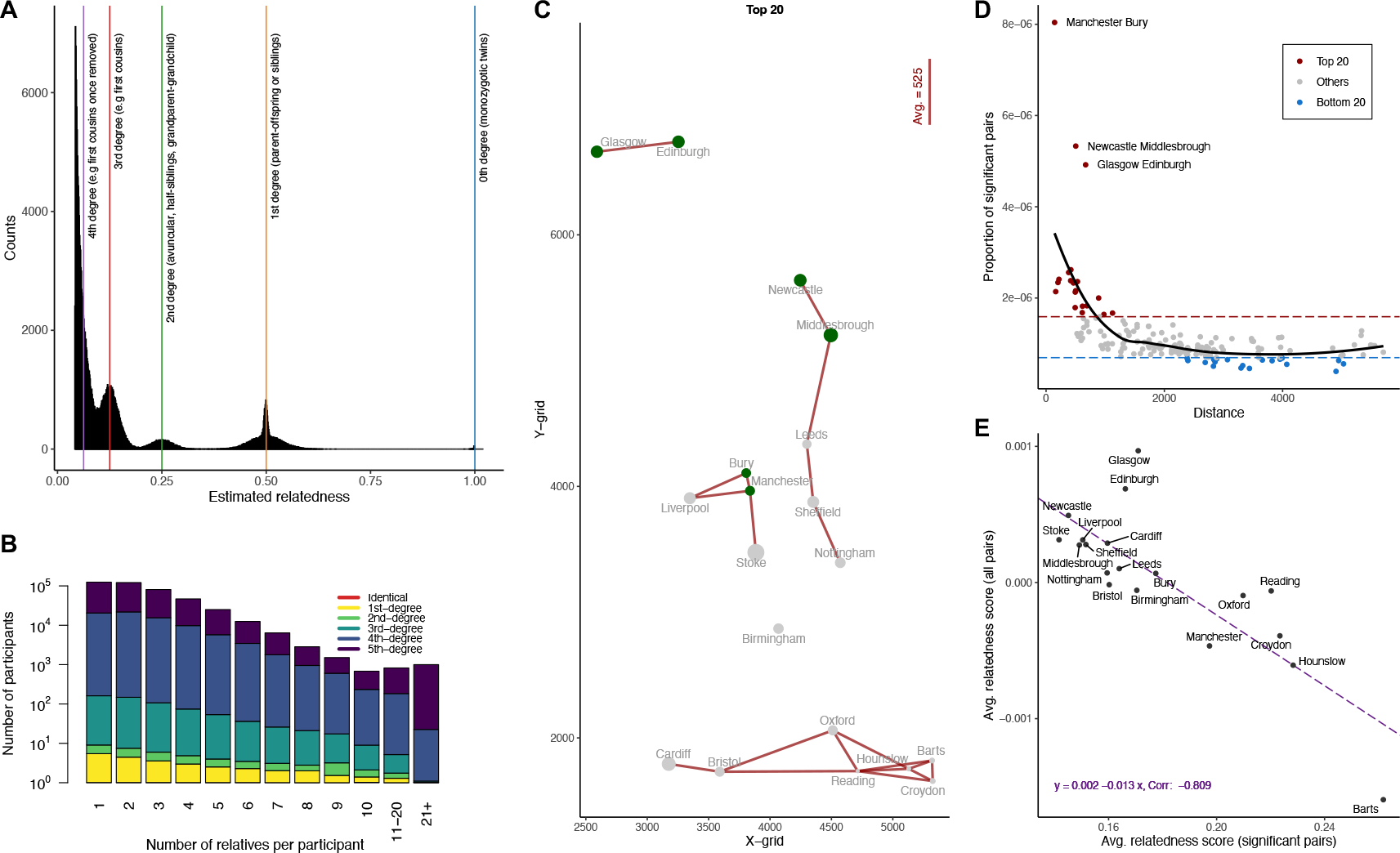
Relatedness in UKB white British ancestry subset (*n*=427,287). (**A**) The histogram shows the distribution of relatedness estimation for 212,120 pair of significant relatedness estimation inferred by deepKin. Colored lines indicate the expectation value of 1.000 (blue), 0.500 (yellow), 0.250 (green), 0.125 (red), and 0.0625 (purple) for zero-, first-, second-, third-, and fourth-degree relationships. (**B**) Distribution of the number of related pairs that participants have in the UKB white British ancestry subset. The height of each bar shows the count of participants (log10 scale) with the stated number of relatives. The colors indicate the proportions of each degree of relatedness within a bar. (**C**) Grid coordinates of 19 assessment centers in UKB and the top 20 pair-wise proportion of cross-cohort significant relatives. The averaging distance is calculated from the average straight-line distance of 20 pairs of cohorts in the plot. The size of the dot indicates the size of the proportion of within-cohort significant relatives. (**D**) The relationship between the proportion of cross-cohort significant relatives and pair-wise distance. (**E**) Regression plot of averaging within-cohort significant relatedness scores (x axis) and overall relatedness scores (y axis) for 19 assessment centers.

We also analyzed the cross-cohort relatedness by assigning participants into 19 cohorts based on their assessment centers. As anticipated, we discovered varying numbers of significant cross-cohort relatives, with a notable pattern: the closer the geographical proximity, the more significant relatives identified (**Figure 5C and 5D**). We define the proportion of cross-cohort significant relatives (*P*) as the ratio between the number of significant cross-cohort relatives (*N*_*sig*_) and the total number of comparisons between the two cohorts (*N*). We observed several cohort pairs with a high proportion of cross-cohort significant relatives, including Manchester and Bury, Glasgow and Edinburgh, and Newcastle and Middlesbrough. The relative proportions of these three pairs were significantly higher than the other top 20 most related pairs. Conversely, certain cohort pairs, such as Glasgow and Cardiff, exhibited a low proportion of cross-cohort significant relatives, likely due to their considerable geographical distance from each other. Specifically, the average grid distances for the top 20 and bottom 20 pairs with the highest and lowest *P* values were 525 and 3,463, respectively (**Figure 5C and Figure S6**).

In each cohort, we considered its within-cohort averaging relatedness score 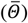 and significant relatedness score 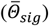 and we observed another notable pattern: the more diverse the population, the closer the relatives are within each cohort (**Table 3 and Figure 5E**). Notably, Glasgow 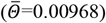 and Edinburgh 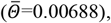, both located in Scotland, exhibited higher levels of relatedness between individuals compared to other cohorts. On the other hand, Barts displayed the lowest averaging relatedness 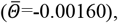, suggesting potential population diversity and even a slightly distinction in population structure (**Figure S7**). Intriguingly, **Figure 5E** revealed a sensible trend: a negative correlation (−0.809) between 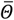 and 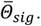. In particular, Barts, with the lowest 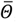, showed the highest 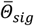 of 0.262, while Glasgow, with the highest 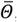, had the lowest 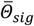 of 0.171. This trend remained significant even after adjusting for sample size or the number of significant pairs within each cohort. These findings robustly indicated both an increase in close relatedness and a potential increase in diversity from north to south across the 19 cohorts.

**Table 3.**
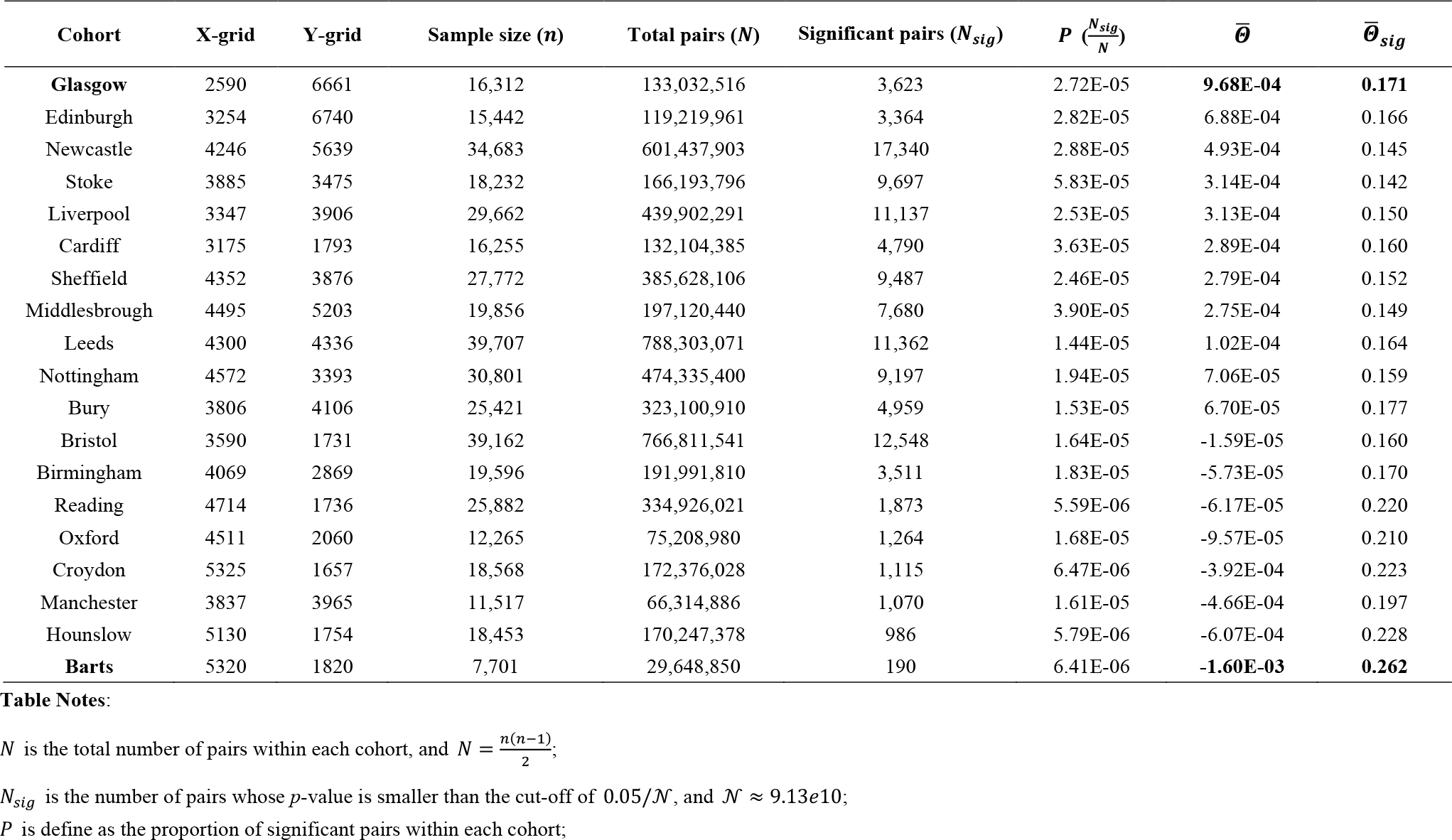
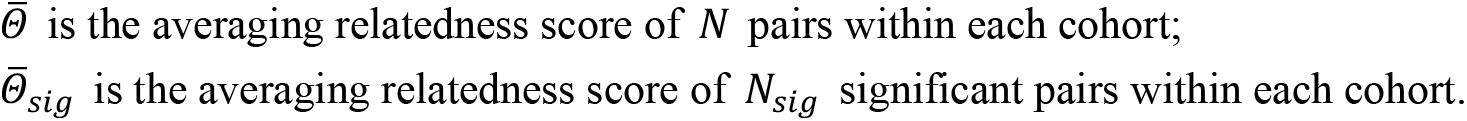
Summary for within-cohort relatedness for 19 UKB assessment centers.

| Cohort | X-grid | Y-grid | Sample size ($n$) | Total pairs ($N$) | Significant pairs ($N_{sig}$) | $P$ ($\frac{N_{sig}}{N}$) | $\bar{\theta}$ | $\bar{\theta}_{sig}$ |
| --- | --- | --- | --- | --- | --- | --- | --- | --- |
| **Glasgow** | 2590 | 6661 | 16,312 | 133,032,516 | 3,623 | 2.72E-05 | **9.68E-04** | **0.171** |
| Edinburgh | 3254 | 6740 | 15,442 | 119,219,961 | 3,364 | 2.82E-05 | 6.88E-04 | 0.166 |
| Newcastle | 4246 | 5639 | 34,683 | 601,437,903 | 17,340 | 2.88E-05 | 4.93E-04 | 0.145 |
| Stoke | 3885 | 3475 | 18,232 | 166,193,796 | 9,697 | 5.83E-05 | 3.14E-04 | 0.142 |
| Liverpool | 3347 | 3906 | 29,662 | 439,902,291 | 11,137 | 2.53E-05 | 3.13E-04 | 0.150 |
| Cardiff | 3175 | 1793 | 16,255 | 132,104,385 | 4,790 | 3.63E-05 | 2.89E-04 | 0.160 |
| Sheffield | 4352 | 3876 | 27,772 | 385,628,106 | 9,487 | 2.46E-05 | 2.79E-04 | 0.152 |
| Middlesbrough | 4495 | 5203 | 19,856 | 197,120,440 | 7,680 | 3.90E-05 | 2.75E-04 | 0.149 |
| Leeds | 4300 | 4336 | 39,707 | 788,303,071 | 11,362 | 1.44E-05 | 1.02E-04 | 0.164 |
| Nottingham | 4572 | 3393 | 30,801 | 474,335,400 | 9,197 | 1.94E-05 | 7.06E-05 | 0.159 |
| Bury | 3806 | 4106 | 25,421 | 323,100,910 | 4,959 | 1.53E-05 | 6.70E-05 | 0.177 |
| Bristol | 3590 | 1731 | 39,162 | 766,811,541 | 12,548 | 1.64E-05 | -1.59E-05 | 0.160 |
| Birmingham | 4069 | 2869 | 19,596 | 191,991,810 | 3,511 | 1.83E-05 | -5.73E-05 | 0.170 |
| Reading | 4714 | 1736 | 25,882 | 334,926,021 | 1,873 | 5.59E-06 | -6.17E-05 | 0.220 |
| Oxford | 4511 | 2060 | 12,265 | 75,208,980 | 1,264 | 1.68E-05 | -9.57E-05 | 0.210 |
| Croydon | 5325 | 1657 | 18,568 | 172,376,028 | 1,115 | 6.47E-06 | -3.92E-04 | 0.223 |
| Manchester | 3837 | 3965 | 11,517 | 66,314,886 | 1,070 | 1.61E-05 | -4.66E-04 | 0.197 |
| Hounslow | 5130 | 1754 | 18,453 | 170,247,378 | 986 | 5.79E-06 | -6.07E-04 | 0.228 |
| **Barts** | 5320 | 1820 | 7,701 | 29,648,850 | 190 | 6.41E-06 | **-1.60E-03** | **0.262** |

## Discussion

Between KING and deepKin, the major difference is that deepKin presents missing elements of KING, such as sampling variance, and consequently sophisticated statistical inference can be conducted. It is known that the resolution of the relationship inference depends on the sampling variance of the estimator. As previously demonstrated for the IBD sharing inference, the sampling variance has been carefully calculated for relatives as defined in a traditional pedigree (Hill and Weir, 2011). However, this general principle has not been fully established in the current framework of genome-wide relatedness measures using SNP data, even though KING has been proposed more than a decade ago, and widely implemented in KING software and other popular GWAS tools such as PLINK. In this study, we present a moment-based estimator deepKin whose sampling variance has been analysed to integrate the characteristics of data, such as MAF and LD, and eventually led to a practically useful asymptotic sampling variance. The availability of the sampling variance brings out a rigorous inference framework for moment-based relatedness estimators. Therefore, deepKin can assess *p*-values for each pair of relatedness, give statistical inference accordingly, divide them as “unrelated” and “related”, and refine classification of each pair into *t*-degree relatives.

Throughout the work, the effective number of markers plays a pivotal role in uncovering the sampling variance of deepKin estimator under the “multiple loci model”. The parameter *m*_*e*_ is a generic population parameter, which characterizes global LD of GWAS markers (Huang *et al*., 2023). Compared to previous studies on the variation of IBD sharing, the counterpart parameter of *m*_*e*_ is the length of genome in terms of Morgan as for linkage-style analysis (Hill and Weir, 2011). Our deepKin guidelines may provide a general solution for choosing optimized SNPs for searching relatedness powerfully, since a larger *m*_*e*_ is more powerful for detecting deeper relatedness. It can be applied in more specific scenarios, such as designing optimized SNP chips for relatedness assessment in forensic applications.

SNP based measures of genome similarity are highly sensitive to the minor allele frequencies in the SNP set. MAFs are influenced by factors such as the choice of SNP genotyping technology and quality control procedures. Although the expected variance under single locus model showed high precision in simulation, it is not practical to assume the same MAF for all variants. At this moment, we extended our derivation to a multiple loci model by assuming a normal distribution of the genotypes after locus-wise standardization, but this assumption may be violated when low-frequency variants are included. We suggest removing low-frequency variants and avoid false positives. Our results demonstrated that the asymptotic variance based on the multiple loci model performed well in both simulations and real dataset applications; future work should focus on extending the assumption of binomial distribution to multiple loci, which takes both the distribution of MAF and LD into consideration.

Although we have demonstrated in principle the theoretical merits and utility of deepKin, there are several questions that should be taken into account for the further development of deepKin. When comparing deepKin with KING, our analysis primarily focused on KING-homo, which performs relatedness estimation and inference within a homogeneous population. deepKin prefers to employ the strategy of selecting non-ancestry-informative (non-AIM) markers to mitigate the potential influence of differentiated allele frequencies. The influence of population stratification on *m*_*e*_, and thus the sampling variance of relatedness estimation is still under investigation. One further investigation is on the union of unilineal relatives and bilineal relatives when detangling the variance of the estimator. In most of the application studies that are based on genome-wide similarity measures, first-degree is usually where they stopped at when distinguishing unilineal relatives (parent-offspring) and bilineal relatives (full siblings). This study, by all means, will be helpful in the construction of a more systematic framework on performing statistical inference on genome-wide similarity measures.

## Supporting information

ExtendData

## Data availability

The genotype and phenotype data of the UK Biobank can be accessed through procedures described on its webpage (https://www.ukbiobank.ac.uk/).

The deepKin estimation and inference have been implemented in an R package “deepKin”. “deepKin” can be downloaded from https://github.com/qixininin/deepKin.

## Acknowledgements

We thank the participants of UK Biobank for making this work possible (application ID 41376). This work was supported by National Natural Science Foundation of China (31771392 to GBC). QXZ thanks the support of the China Scholarship Council program (project ID: 202306320498).

## Appendix I Variance of deepKin under the single locus model

We derived the variance of deepKin to the context of binomial distribution. If we consider two diploid individuals, each with two alleles (*A* and *a*) at one locus. The allele frequencies are *p* and *q* for allele *A* and *a*, respectively. For a pair of relatives, the probabilities of two alleles conditioning on the probability of relatedness *θ* is:

| Allelic pair for relatives | Probability |
| --- | --- |
| $A/A$ | $\theta p + (1 - \theta)p^2$ |
| $A/a$ | $(1 - \theta)pq$ |
| $a/A$ | $(1 - \theta)pq$ |
| $a/a$ | $\theta q + (1 - \theta)q^2$ |

The above table can be applied to their respect second allele pair. The probabilities of sixteen genotype combinations between two individuals are filled in this 4 × 4 table, which is symmetric:

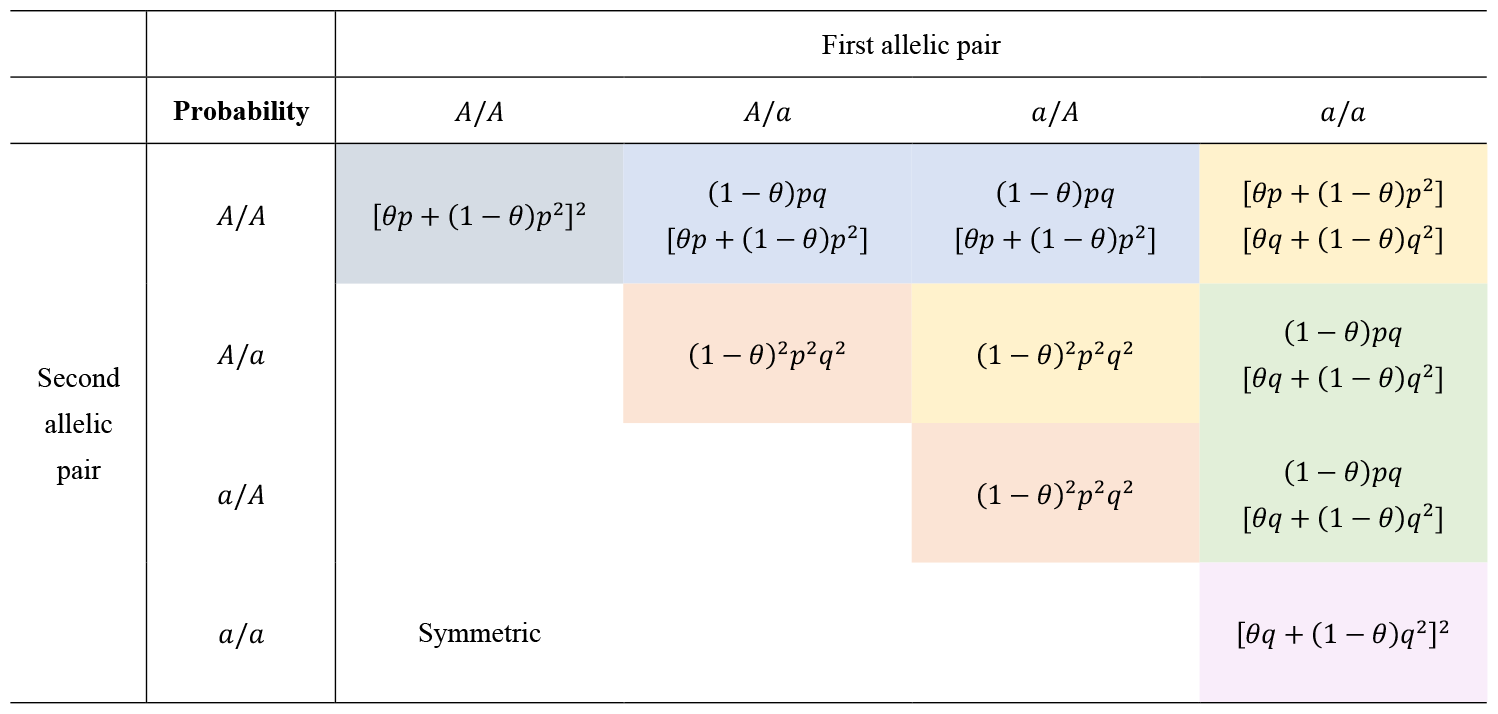

|  |  | First allelic pair |  |  |  |
| --- | --- | --- | --- | --- | --- |
| | | $A/A$ | $A/a$ | $a/A$ | $a/a$ |
| Second allelic pair | $A/A$ | $[\theta p + (1 - \theta)p^2]^2$ | $(1 - \theta)pq$<br>$[\theta p + (1 - \theta)p^2]$ | $(1 - \theta)pq$<br>$[\theta p + (1 - \theta)p^2]$ | $[\theta p + (1 - \theta)p^2]$<br>$[\theta q + (1 - \theta)q^2]$ |
| | $A/a$ | | $(1 - \theta)^2 p^2 q^2$ | $(1 - \theta)^2 p^2 q^2$ | $(1 - \theta)pq$<br>$[\theta q + (1 - \theta)q^2]$ |
| | $a/A$ | | | $(1 - \theta)^2 p^2 q^2$ | $(1 - \theta)pq$<br>$[\theta q + (1 - \theta)q^2]$ |
| | $a/a$ | Symmetric | | | $[\theta q + (1 - \theta)q^2]^2$ |

Ignoring the order of two individuals, we can recalculate the frequencies by summing up the corresponding probabilities in each cell. As individual genotypes are standardized by 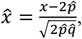, where *x* is coded as the number of minor alleles. deepKin score between individual *i* and individual *j* is estimated by 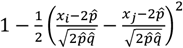. 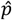 and 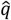 are the estimated allele frequencies of *A* and *a*. The rearranged probabilities and relatedness scores are:

| Genotype pair | Genotype code | Probability ($f_i$) | Relatedness ($X_i$) |
| --- | --- | --- | --- |
| $\{AA, AA\}$ | $\{2, 2\}$ | $[\theta p + (1 - \theta)p^2]^2$ | 1 |
| $\{Aa, Aa\}$ | $\{1, 1\}$ | $4(1 - \theta)^2 p^2 q^2 + 2\theta pq$ | 1 |
| $\{aa, aa\}$ | $\{0, 0\}$ | $[\theta q + (1 - \theta)q^2]^2$ | 1 |
| $\{AA, Aa\}$ | $\{2, 1\}$ | $4\theta(1 - \theta)p^2 q + 4(1 - \theta)^2 p^3 q$ | $\frac{4pq - 1}{4pq}$ |
| $\{Aa, aa\}$ | $\{1, 0\}$ | $4\theta(1 - \theta)pq^2 + 4(1 - \theta)^2 pq^3$ | $\frac{4pq - 1}{4pq}$ |
| $\{AA, aa\}$ | $\{2, 0\}$ | $2(1 - \theta)^2 p^2 q^2$ | $\frac{pq - 1}{pq}$ |

Expectation

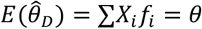

Variance

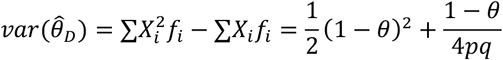

## Appendix II Variance of deepKin under multiple loci model

The expectation and variance for deepKin 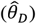 are,

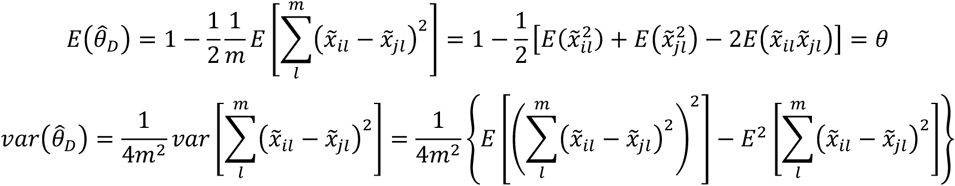

The first term of *var*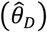 can be decomposed as

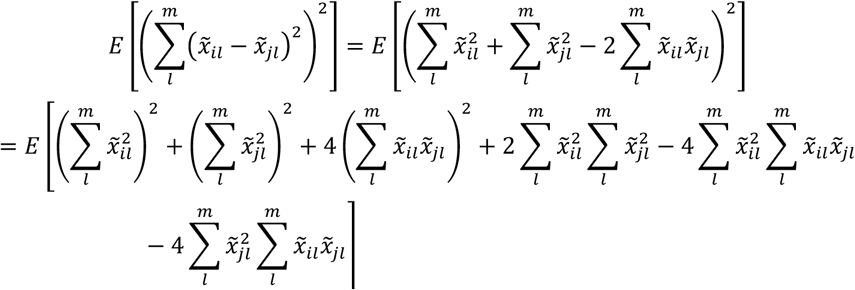

| Term | Expectation |
| --- | --- |
| $E[(\sum_l^m \tilde{x}_{il}^2)^2] \text{ or } E[(\sum_l^m \tilde{x}_{jl}^2)^2]$ | $m^2 + 2m + 2 \sum_l^m \sum_{l' \neq l}^m \rho_{ll'}^2$ |
| $E \left[ \left( \sum_l^m \tilde{x}_{il}\tilde{x}_{jl} \right)^2 \right]$ | $m + m\theta^2 + m^2\theta^2 + (1 + \theta^2) \sum_l^m \sum_{l' \neq l}^m \rho_{ll'}^2$ |
| $E \left( \sum_l^m \tilde{x}_{il}^2 \sum_l^m \tilde{x}_{jl}^2 \right)$ | $2m\theta^2 + m^2 + 2\theta^2 \sum_l^m \sum_{l' \neq l}^m \rho_{ll'}^2$ |
| $E(\sum_l^m \tilde{x}_{il}^2 \sum_l^m \tilde{x}_{il}\tilde{x}_{jl}) \text{ or } E(\sum_l^m \tilde{x}_{jl}^2 \sum_l^m \tilde{x}_{il}\tilde{x}_{jl})$ | $2m\theta + m^2\theta + 2\theta \sum_l^m \sum_{l' \neq l}^m \rho_{ll'}^2$ |

The second term of 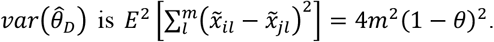.

Therefore,

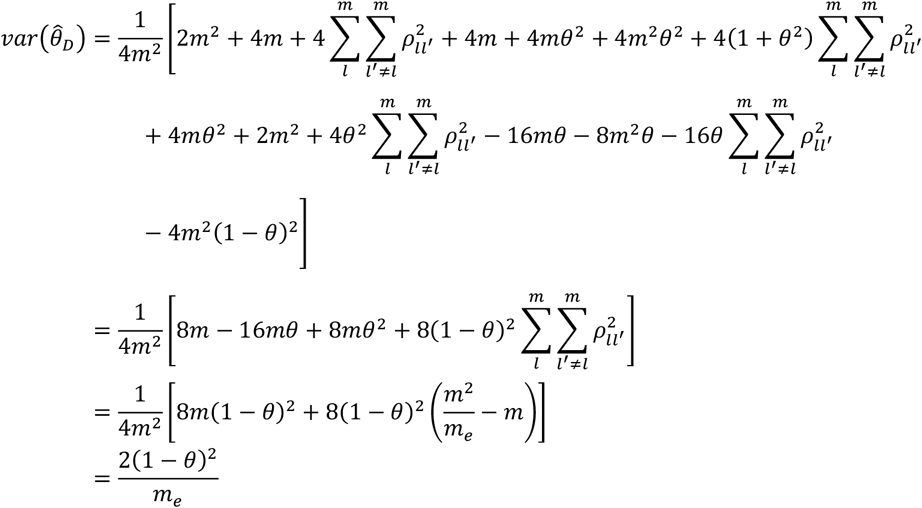

## Appendix III

Let *Z*_*t*_ = *Z*_*t*+1_, When *t* < *x* < *t* + 1

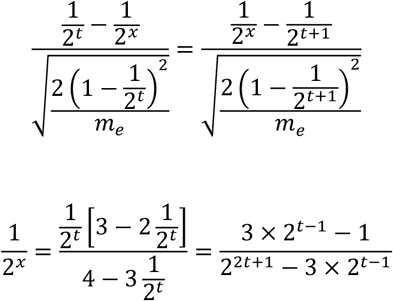

| Degree | Geometric mean-based lower boundary | DeepKin variance-based lower boundary |
| --- | --- | --- |
| 0 (0.1) | 0.707 | 0.881 |
| 1 | 0.354 | 0.400 |
| 2 | 0.177 | 0.192 |
| 3 | 0.088 | 0.095 |
| 4 | 0.044 | 0.047 |
| 5 | 0.022 | 0.024 |
| ... | ... |  |
| $t$ | $\frac{1}{2^{(2t+1)/2}}$ | $\frac{3 \times 2^{t-1} - 1}{2^{2t+1} - 3 \times 2^{t-1}}$ |

When *t* = 0, the sampling variance becomes 0, but we cannot use threshold =1 to separate first-degree and identical pairs. Therefore, when *x* is between 0 and 1, we set *t*=0.1, and the corresponding threshold becomes 0.881.

## Appendix IV

For hypothesis on a statistic *T*, which follows 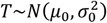 for *H*_0_ and follows 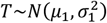 for *H*_1_. We assume that 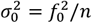 and 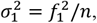, where *n* is the number of sample size. For one-tailed hypothesis that *H*_0_: *μ* = *μ*_0_, and *H*_1_: *μ* ≠ *μ*_0_, the minimal number of *n* is,

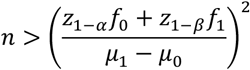

In this case, *H*_0_ : *μ*_0_ = *θ* = 0, and *H*_1_ : *μ*_1_ = *θ* > 0. Given 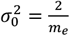 and 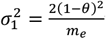 the minimum number of *m*_*e*_ is,

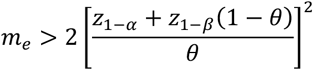

## Supplementary tables and figures

**Figure S1.**
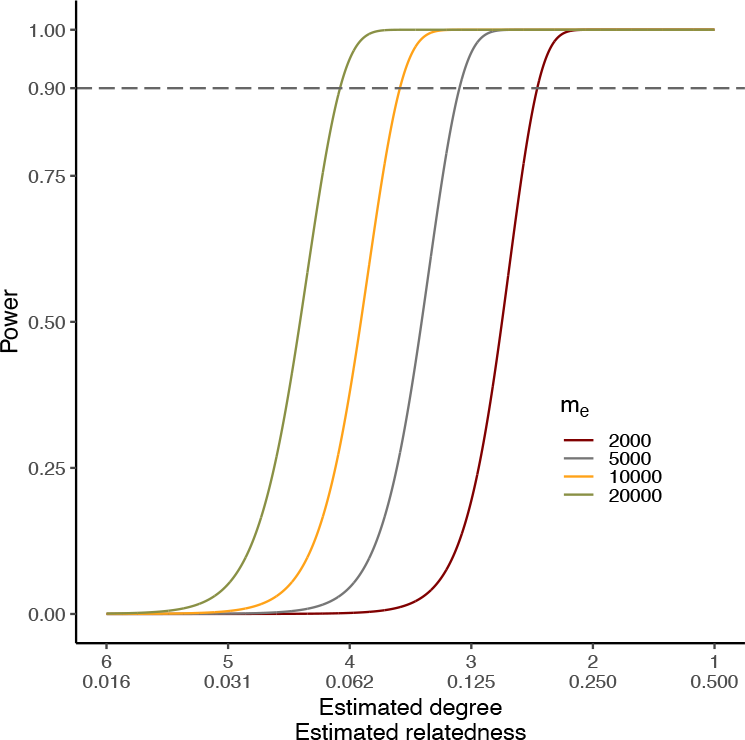
The power of detecting each degree of relatedness calculated based on **Eq 7** under four different effective number of markers (*m*_*e*_=2,000, 5,000, 10,000, and 2,000). The dashed grey line indicates 90% power. A range of degrees from zero to six are considered. Suppose Type I error rate of *α* = 0.05/*N* and Type II error rate of *β*=0.1, where *N* = 40,000.

**Figure S2.**
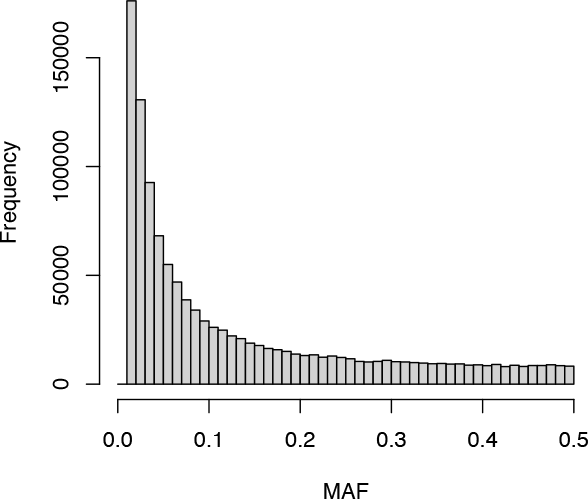
The histogram of MAF distribution in Oxford 3K demo after quality control (MAF > 0.01; HWE test *p*-value > 1e-7; no locus missingness).

**Figure S3.**
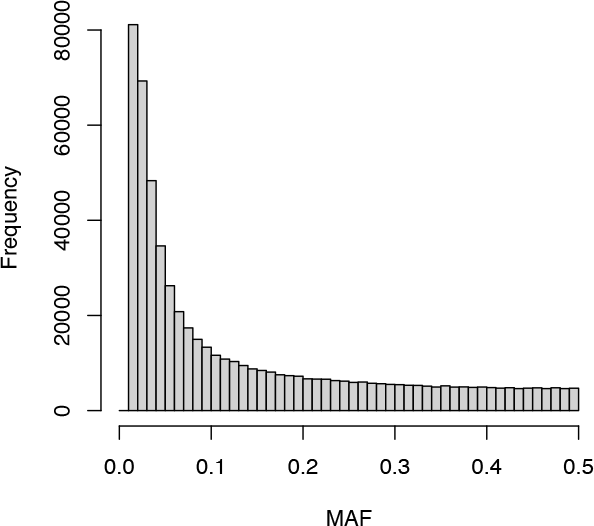
The histogram of MAF distribution for UKB white British ancestry subset after quality control (MAF > 0.01; HWE test *p*-value > 1e-7; no locus missingness).

**Figure S4.**
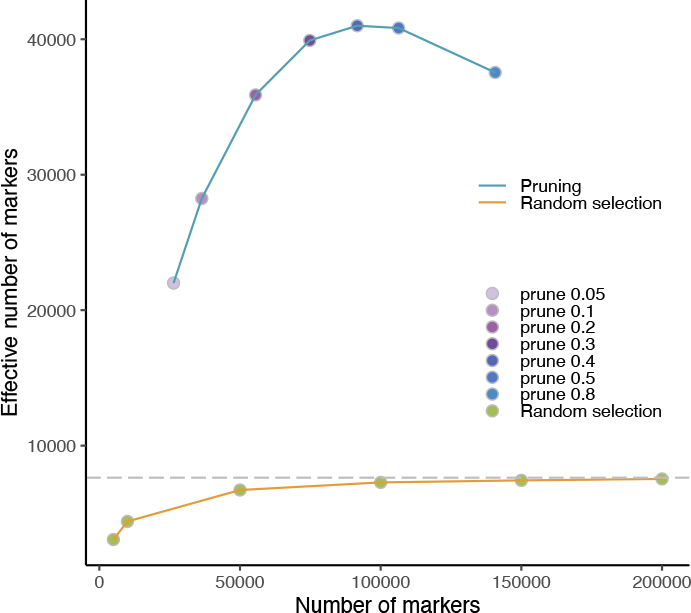
The distribution of effective number of markers under 13 circumstances, 6 of which were random selected markers but of different numbers (*m* = 5,000, 10,000, 50,000, 100,000, 150,000, and 200,000) and 7 were with different pruning thresholds (*r*^2^< 0.05, 0.1, 0.2, 0.3, 0.4, 0.5, and 0.8). The grey dashed line indicated the *m*_*e*_ of all *m* = 298,211 variants.

**Figure S5.**
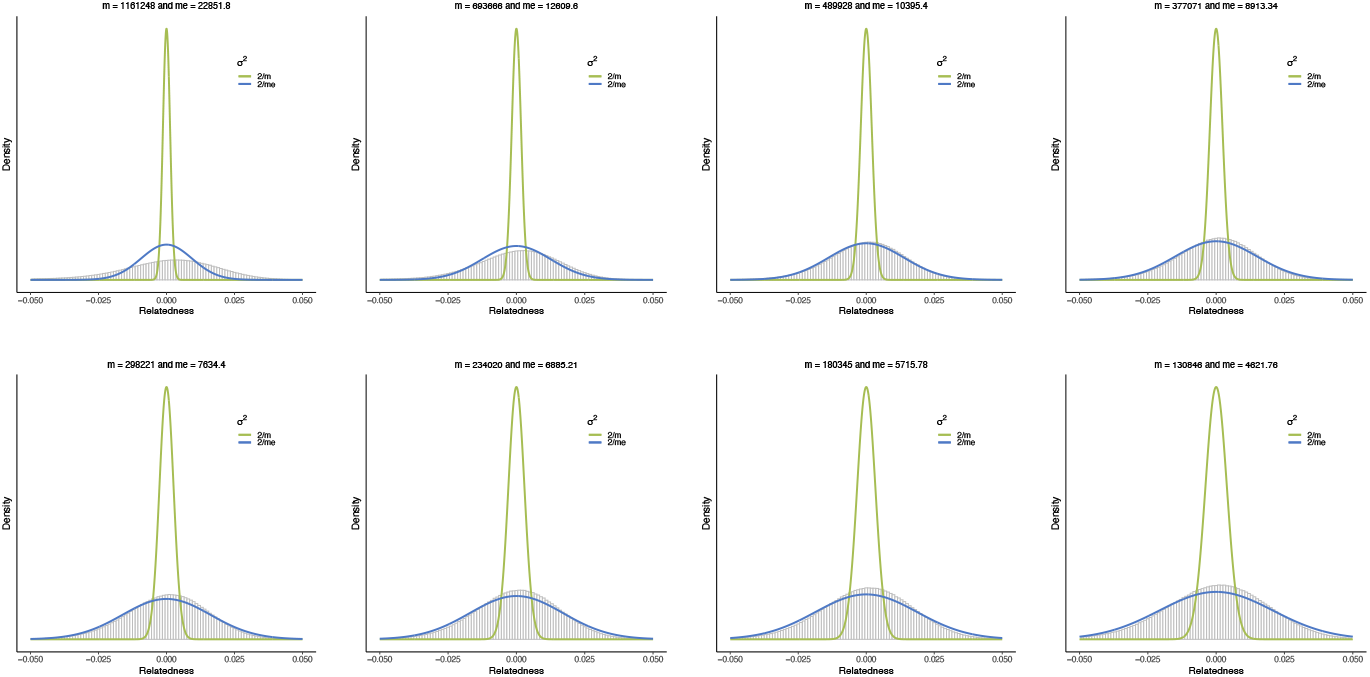
The histogram of estimated relatedness scores based on different MAF thresholds. MAF thresholds from left to right, from up to bottom are 0.01, 0.05, 0.10, 0.15, 0.20, 0.25, 0.30, and 0.35. The green and blue curves indicate the normal distribution of *N*(0, 2⁄*m*) and *N*(0, 2⁄*m*_*e*_), respectively.

**Figure S6.**
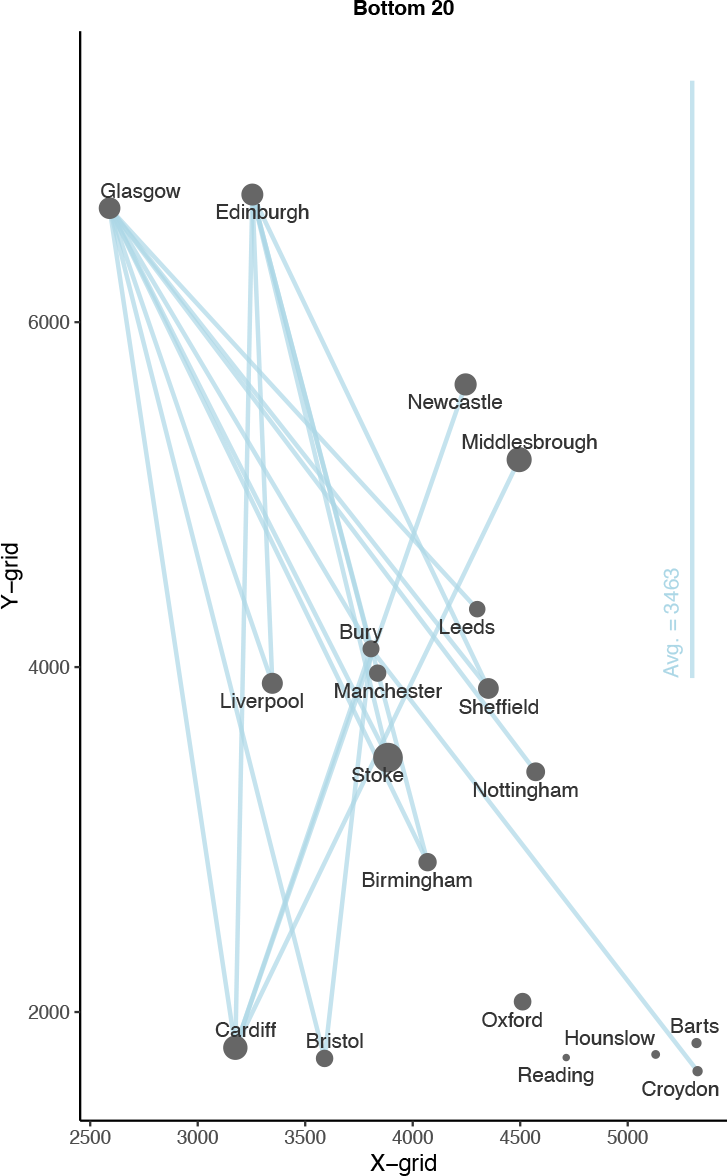
Grid coordinates of 19 assessment centers in UKB and the bottom 20 pair-wise proportion of cross-cohort significant relatives. The averaging distance is calculated from the average straight-line distance of 20 pairs of cohorts in the plot. The size of the dot indicates the size of the proportion of within-cohort significant relatives.

**Figure S7.**
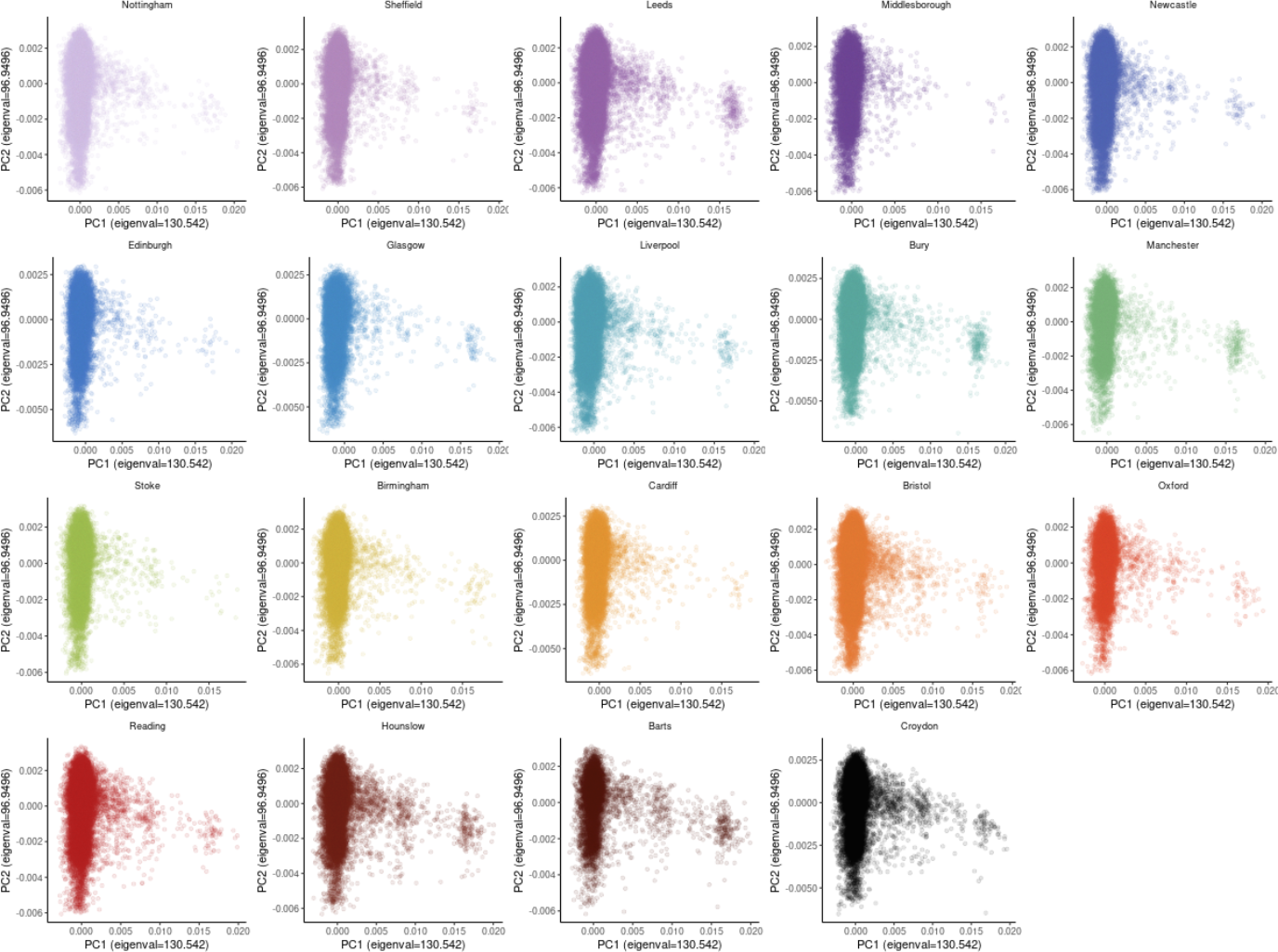
The distribution of the first two PCs for individuals in the 19 UKB cohorts.

**Table S1.**
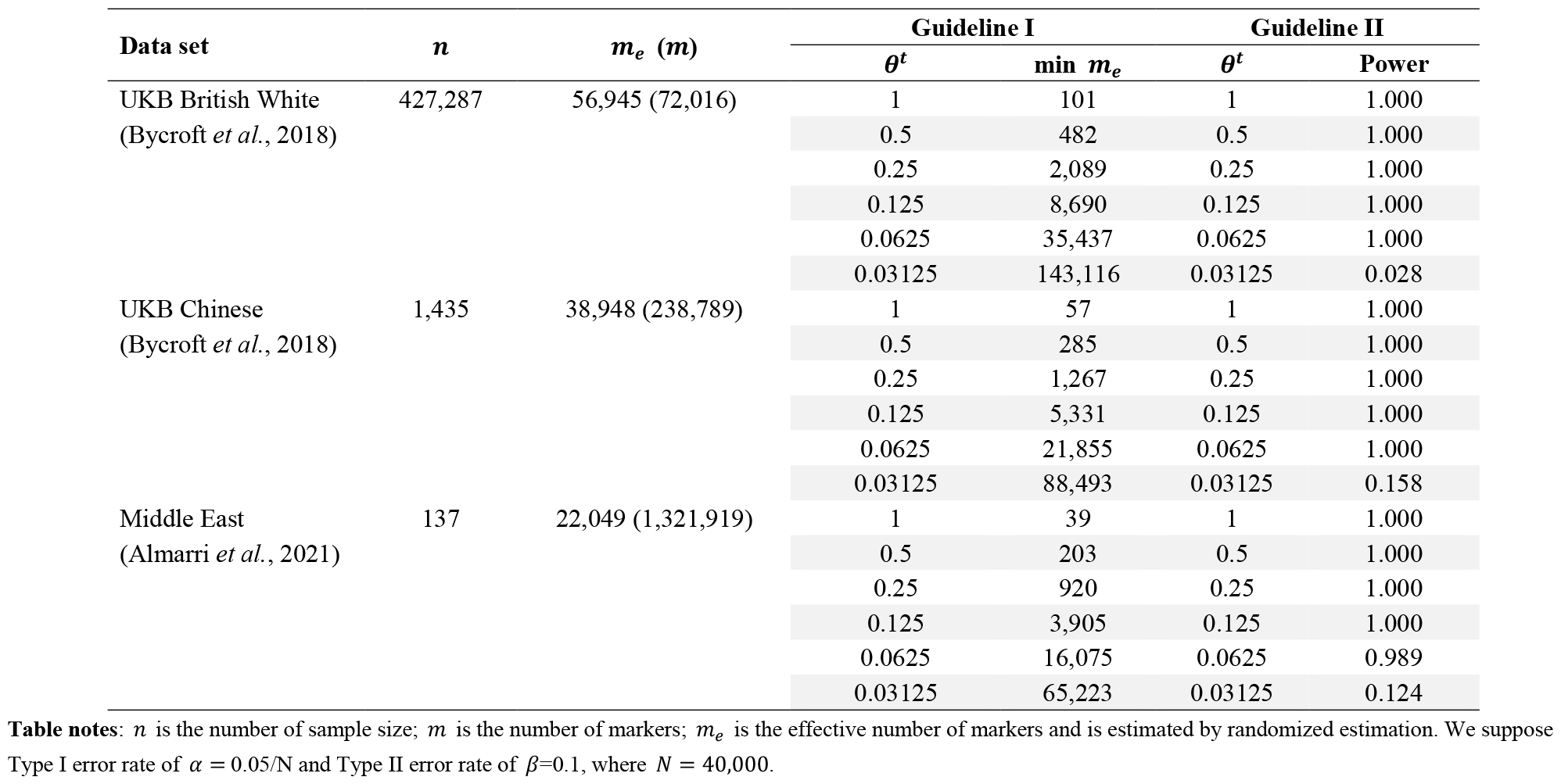
Real dataset examples for applying the deepKin guidelines.

| Data set | $n$ | $m_e$ ($m$) | Guideline I | | Guideline II | |
| --- | --- | --- | --- | --- | --- | --- |
| | | | $\theta^t$ | $\min m_e$ | $\theta^t$ | Power |
| UKB British White<br>(Bycroft *et al.*, 2018) | 427,287 | 56,945 (72,016) | 1 | 101 | 1 | 1.000 |
|  |  |  | 0.5 | 482 | 0.5 | 1.000 |
|  |  |  | 0.25 | 2,089 | 0.25 | 1.000 |
|  |  |  | 0.125 | 8,690 | 0.125 | 1.000 |
|  |  |  | 0.0625 | 35,437 | 0.0625 | 1.000 |
|  |  |  | 0.03125 | 143,116 | 0.03125 | 0.028 |
| UKB Chinese<br>(Bycroft *et al.*, 2018) | 1,435 | 38,948 (238,789) | 1 | 57 | 1 | 1.000 |
|  |  |  | 0.5 | 285 | 0.5 | 1.000 |
|  |  |  | 0.25 | 1,267 | 0.25 | 1.000 |
|  |  |  | 0.125 | 5,331 | 0.125 | 1.000 |
|  |  |  | 0.0625 | 21,855 | 0.0625 | 1.000 |
|  |  |  | 0.03125 | 88,493 | 0.03125 | 0.158 |
| Middle East<br>(Almarri *et al.*, 2021) | 137 | 22,049 (1,321,919) | 1 | 39 | 1 | 1.000 |
|  |  |  | 0.5 | 203 | 0.5 | 1.000 |
|  |  |  | 0.25 | 920 | 0.25 | 1.000 |
|  |  |  | 0.125 | 3,905 | 0.125 | 1.000 |
|  |  |  | 0.0625 | 16,075 | 0.0625 | 0.989 |
|  |  |  | 0.03125 | 65,223 | 0.03125 | 0.124 |

**Table notes:** $n$ is the number of sample size; $m$ is the number of markers; $m_e$ is the effective number of markers and is estimated by randomized estimation. We suppose Type I error rate of $\alpha = 0.05/N$ and Type II error rate of $\beta=0.1$ , where $N = 40,000$ .

**Table S2.** QC details for four SNP sets and the numbers of pairs exceed relatedness cut-offs based on two normal GRM cutoffs in the Oxford demo.

| SNP set | QC details | Number of SNPs | $m_e$ | $\delta$ | GRM Cut-off | Number of pairs |
| --- | --- | --- | --- | --- | --- | --- |
| Set 1 | MAF>0.05 | 693,666 | 12,610 | 3.827 | $\theta>0.05$ | 2,381 |
| | | | | | $\theta>0.025$ | 47,268 |
| Set 2 | MAF>0.2 | 298,221 | 7,634 | 3.465 | $\theta>0.05$ | 539 |
| | | | | | $\theta>0.025$ | 72,032 |
| Set 3 | MAF>0.2;<br>eigenGWAS p-value>0.05 | 237,642 | 10,424 | 3.690 | $\theta>0.05$ | 108 |
| | | | | | $\theta>0.025$ | 22,564 |
| Set 4 | MAF>0.2;<br>LD pruning (50 5 0.1) | 36,425 | 28,244 | 4.409 | $\theta>0.05$ | 60 |
| | | | | | $\theta>0.025$ | 655 |

**Table Note:** 3,000 individuals and a total number of ~4.5 million comparisons are considered.
\* $\delta$ is the deepest significant degree that the data would support to detect from unrelated individuals.
\* $m_e$ is estimated by randomized estimation.

**Table S3.** Summary of the significant related pairs in UKB British white dataset (*α* = 0.05/*N, N* = 9.13*e*10).

| Degree | Identical | 1st-degree | 2nd-degree | 3rd-degree | 4th-degree | 5th-degree | Total |
| --- | --- | --- | --- | --- | --- | --- | --- |
| Number of pairs | 162 | 25,699 | 9,455 | 53,221 | 91,977 | 31,606 | 212,120 |

